# Male host suppressors mask *Wolbachia*-induced cytoplasmic incompatibility in the invasive pest *Drosophila suzukii*

**DOI:** 10.64898/2026.11.12.766142

**Authors:** Svitlana Serga, Mountassira Matin, Chooi Lin Phooi, Candice Deschamps, Lily Cesari, Benjamin Kaltenbach, Emilie Mendes, Anne Loiseau, Caroline Tatard, Laure Sauné, Sabine Nidelet, Laure Benoit, Marie Lapègue, Hervé Colinet, Arnaud Estoup, Benjamin Loppin, Mathieu Gautier, Nicolas O. Rode

**Affiliations:** CBGP, Univ Montpellier, CIRAD, INRAE, IRD, Institut Agro Montpellier, Montpellier, France; V.N. Karazin Kharkiv National University, 61022, Kharkiv, Ukraine; University of Rennes, CNRS, ECOBIO [(Ecosystèmes, biodiversité, évolution)] - UMR 6553, 35042, Rennes, France; Laboratoire de Biologie et Modélisation de la Cellule, École Normale Supérieure de Lyon, CNRS UMR5239, Lyon 69007, France

**Keywords:** insect pest, invasive species, male sterility, reproductive manipulation, host-symbiont interaction

## Abstract

Cytoplasmic incompatibility (CI) is the most widespread reproductive manipulation induced by the endosymbiotic bacteria *Wolbachia* in arthropods. When infected males mate with uninfected females, some or all embryos die, which favors infected females and drives the spread of the symbiont through host populations. The *Wolbachia cif* genes that cause CI have been identified, but they do not explain why CI strength varies among host genotypes. The invasive fruit pest *Drosophila suzukii* illustrates a related paradox: its *Wolbachia* strain *w*Suz carries *cif* genes, yet CI has not been detected in this host species. Combining experimental crosses with genomic and cytological approaches, we show that *w*Suz does induce CI, with paternal chromosomes showing segregation defects during the first zygotic division. However, CI strength ranges from undetectable to 67% among males from four European host lines. This variation was explained neither by *Wolbachia* titer in testes nor by the *Wolbachia* genome, which was nearly identical among isolates and invariant at *cif* loci. Instead, CI strength depended on the genotype of infected males, whereas the genotype of uninfected females had no detectable effect. Reciprocal F1 and F2 crosses revealed that CI is masked by partially dominant autosomal host suppressors, probably involving a few loci. Masking by host suppressors thus explains the apparent absence of CI in *D. suzukii*, and host suppressors of CI may also shape the efficacy and evolutionary success of *Wolbachia*-based pest and vector control strategies.

**Significance Statement:** Wolbachia bacteria infect about half of arthropod species and spread by manipulating host reproduction: when infected males mate with uninfected females, some or all of the embryos die. This cytoplasmic incompatibility is used to suppress mosquito populations and is being developed against the invasive fruit pest *Drosophila suzukii*. Yet, although the native *Wolbachia* of *D. suzukii* carries the genes that cause incompatibility, it has not been detected. We show that the bacterium does induce it, but that partially dominant host suppressors acting in males mask it to varying degrees, whereas the bacterial genome is virtually identical across host lines. The success of *Wolbachia*-based pest and vector control may therefore depend on host genetics, not only on the *Wolbachia* strain.

## Introduction

Maternally inherited endosymbionts that alter the reproduction of invertebrates represent potent drivers of host evolution (Werren et al., 2008). Among the roughly 50% of arthropod species infected by the endosymbiotic bacteria *Wolbachia*, the most common alteration of reproduction is cytoplasmic incompatibility (CI; Turelli et al., 2022). This form of conditional male sterility occurs when crosses between infected males and uninfected females result in the death of some or all embryos, depending on the strength of CI (Werren et al., 2008). Embryonic lethality is prevented when the female carries the same or a compatible *Wolbachia* strain, a process known as rescue. Because infected females are compatible with both infected and uninfected males, whereas uninfected females are compatible only with uninfected males, CI gives infected females a reproductive advantage. This allows *Wolbachia* to spread once it exceeds a threshold frequency, provided the advantage outweighs the costs of infection and imperfect maternal transmission (Turelli et al., 2022).

CI in *Wolbachia* is caused by two genes, *cifA* and *cifB* (cytoplasmic incompatibility factors), which occur as adjacent pairs, generally within prophage WO regions (Beckmann et al., 2017; LePage et al., 2017). A single *Wolbachia* genome can carry several *cifA-cifB* pairs belonging to distinct phylogenetic types (Martinez et al., 2021). CifA and CifB are thought to act as a toxin-antidote system, in which sperm carrying the CifB toxin triggers embryonic lethality unless the egg is infected with a compatible *Wolbachia* strain expressing CifA (Beckmann et al., 2019; Hochstrasser, 2023).

While environmental factors such as *Wolbachia* titer, male age, larval rearing density, or temperature can modulate the expression of CI, accumulating evidence indicates that genetic variation, both in the symbiont and the host, plays a central role in determining CI strength (Shropshire et al., 2020; Turelli et al., 2022). Interestingly, although intact *cifA–cifB* pairs are strongly associated with CI, their presence does not fully explain the substantial variation in CI expression observed across host–*Wolbachia* associations. For example, several *Wolbachia* strains harbor at least one presumably functional pair of *cifA and cifB* genes, but show low or undetectable CI (Martinez et al., 2021). Such cases are generally attributed to the symbiont, yet transfers of *Wolbachia* from native to novel host species have shown that the host nuclear background can modulate CI strength (Bordenstein et al., 2003; Martinez et al., 2015; Poinsot et al., 1998; Walker et al., 2011; Zabalou et al., 2008). Such artificial host-symbiont combinations, however, reveal little about the variation that occurs within natural populations and on which selection acts. Theory predicts that host alleles reducing CI in males should be favored, because infected males that carry them sire viable offspring when they mate with uninfected females (Kauai & Wybouw, 2025; Koehncke et al., 2009; Turelli, 1994). Consistent with this prediction, CI strength varies with male host genotype within several species (Cooper et al., 2017; Numajiri et al., 2025; Reynolds & Hoffmann, 2002; Wybouw et al., 2022). Yet, the genetic architecture of host suppression remains uncharacterized: crossing experiments suggest a polygenic basis in the spider mite *Tetranychus urticae* (Wybouw et al., 2022) and the bean weevil *Callosobruchus analis* (Numajiri et al., 2025), but the level of dominance and chromosomal location of suppressor loci remain unknown. Addressing this gap is critical for both advancing our understanding of host-symbiont coevolution and improving *Wolbachia*-based biocontrol strategies.

The invasive fruit pest *Drosophila suzukii* provides a striking example of this decoupling between intact *cif* genes and CI expression. Its native *Wolbachia* strain, *w*Suz, carries *cif* genes (Conner et al., 2017), yet crossing experiments have so far detected no CI (Cattel, Kaur, et al., 2016; Hamm et al., 2014). This apparent paradox makes the *D. suzukii*-*w*Suz association a well-suited system to investigate how host and symbiont genotypes shape CI strength within a species. Moreover, *w*Suz persists at intermediate and regionally variable frequencies in natural populations (Cattel, Kaur, et al., 2016; Hamm et al., 2014), the situation in which theory predicts the strongest selection on host suppressors. Understanding how *Wolbachia* and host genetic backgrounds determine CI expression is critical, as CI has been proposed as a basis for incompatible insect technique (IIT) approaches to suppress *D. suzukii* populations (Nikolouli et al., 2018). Here, we investigate the respective roles of *Wolbachia*, paternal and maternal host genetic factors in driving CI in *D. suzukii* using experimental crosses in combination with genomic and cytological approaches. Our results reveal substantial intraspecific variation in CI strength, ranging from no detectable incompatibility to strong (67%) embryonic lethality. Our findings indicate that phenotypic variation is driven by male genotypes and more specifically by partially dominant autosomal suppressors carried by males, and not by differences among *Wolbachia* genomes, among maternal genotypes or in *Wolbachia* titer.

## Results

### *w*Suz induces bona fide CI whose strength varies among host lines

For each of four *Wolbachia*-infected European lines of *D. suzukii* (Balandran [Bal], Trento, Rennes and Paris [Par6]; *SI Appendix*, Table S1), we compared the hatch rate of a test cross between uninfected females and infected males (where CI could occur) with that of three control crosses: uninfected females × uninfected males, infected females × uninfected males, and infected females × infected males (Fig. 1A). While we could not detect any hatch rate reduction in test crosses with Rennes and Par6 males, we detected strong reductions in hatch rates in test crosses with Bal males compared to all control crosses (67% relative control cross 1; 95% confidence interval: 58%-75%) and to a lesser extent for Trento males (47%, 39%-55%) (Fig. 1B, *SI Appendix*, Table S2). Two lines of evidence show that this reduction in hatch rate results from CI. First, it depended on *Wolbachia*: it was no longer observed in test crosses with Bal and Trento males cured of *Wolbachia* using tetracycline (*SI Appendix*, Fig. S1, Table S3). Second, it was rescued by maternal infection: hatch rate was not reduced when infected males were crossed with infected females from the same line (Fig. 1B, Table S2). Together, these results demonstrate that *w*Suz induces *bona fide* CI in at least two *D. suzukii* lines.

**Fig. 1.**
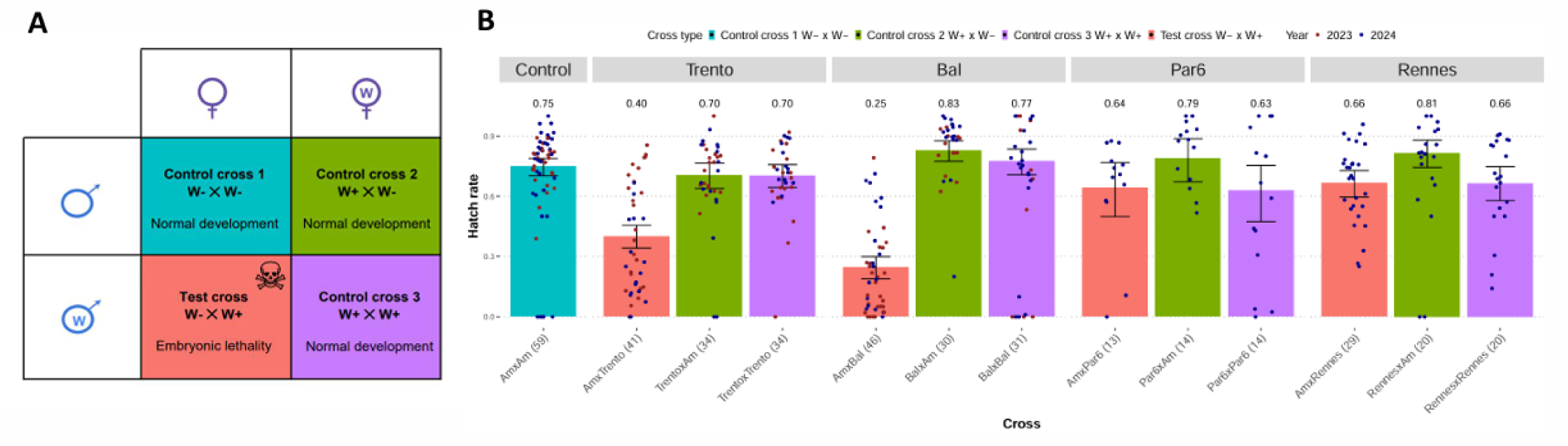
Cytoplasmic incompatibility assay for four European *D. suzukii* lines infected with *Wolbachia.* (A) Four types of crosses used to measure cytoplasmic incompatibility. In the presence of CI, hatch rates in test crosses (W− female × W+ male) are significantly lower than in control crosses (W− × W−; W+ × W−and W+ × W+). (B) Observed hatch rate and fitted hatch rate excluding structural zeros in test and control crosses. Hatch rates in test crosses with either Bal or Trento males were consistently lower in test crosses than in control crosses. This was not the case for test crosses with Par6 or Rennes males. The four types of crosses are indicated as “female line × male line” (W+ : infected line, W-: uninfected line). For each cross, the sample size of the number of pairs used to fit the model is indicated below each bar in parentheses. Model selection was performed using AICc. Error bars represent 95%-confidence intervals calculated using parametric bootstrap with 500 iterations (*SI Appendix*, Table S2).

To test whether *w*Suz-induced embryonic lethality shows the cytological hallmarks of CI described in other *Drosophila* species, we examined early embryos from a test cross (uninfected Am females × infected Bal males) and from a rescue cross (infected Bal females × infected Bal males; Fig. 2). We observed that some embryos showed chromosome segregation defects during early nuclear divisions (Fig. 2A), whereas others went through the first mitosis normally (Fig. 2B). In particular, we observed embryos in nuclear cycle 1 with the two daughter nuclei connected by a chromatin bridge (Fig. 2A). This phenotype, which is typical of CI, results from delayed segregation of paternal chromosomes at the first mitosis. Later embryos showed various phenotypes, ranging from severe aneuploid nuclear divisions to an absence of visible defects (Fig. 2C,E). These defects were not observed in embryos from a rescue cross between infected Bal individuals (Fig. 2G,H).

**Fig. 2.**
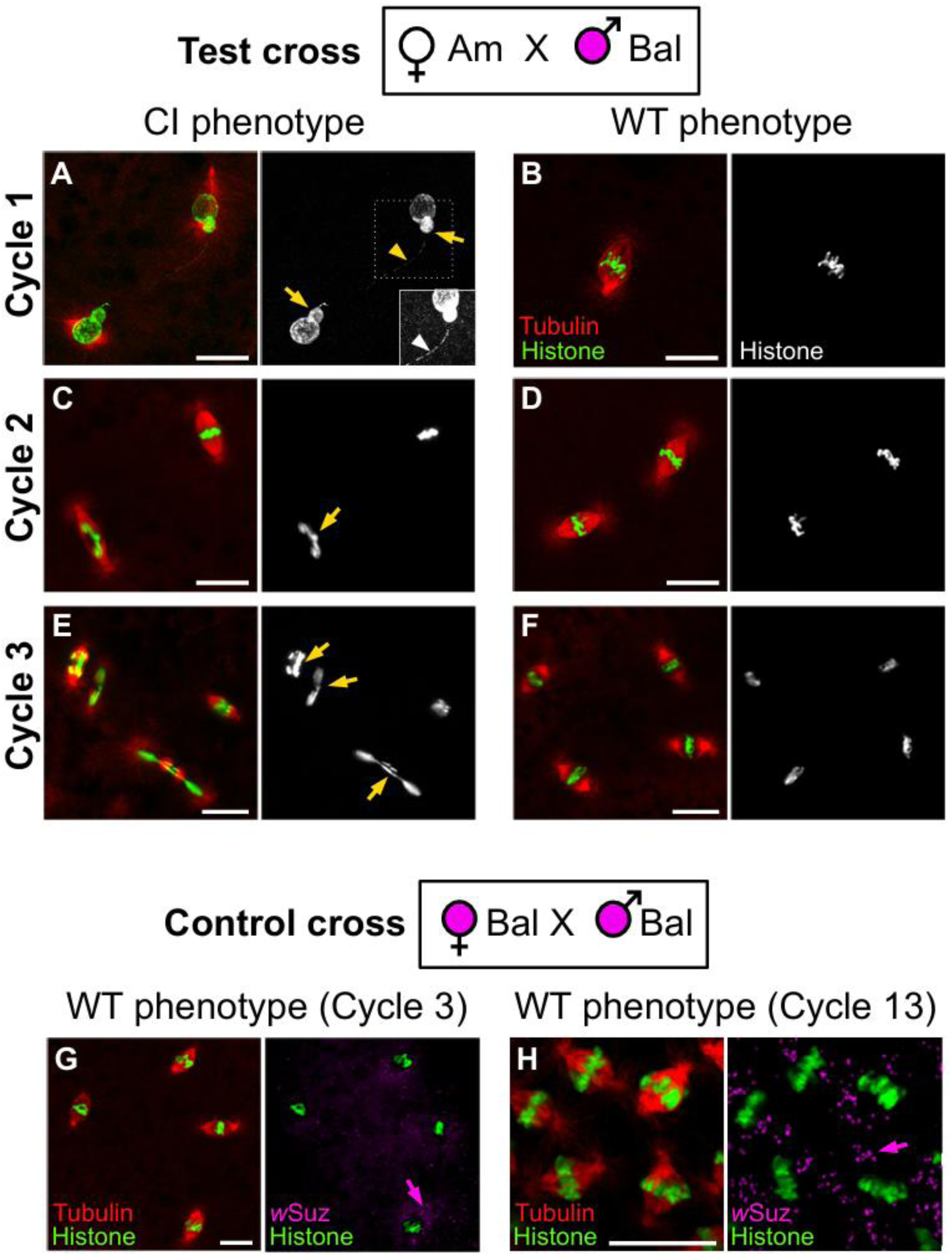
Cytological analyses of *w*Suz CI (Test cross) and rescue (Control cross 3) in *D. suzukii* embryos. (A-F) Confocal images of early *D. suzukii* embryos from a test cross between infected Bal males and uninfected Am females. Chromosomes are labeled with anti-pan histone (green) and mitotic spindles are labeled with anti-alpha-tubulin (red). (A) End of nuclear cycle 1 showing a typical CI phenotype. Note the thin chromatin bridge (arrowheads) connecting the late segregating chromosomes of presumably paternal origin (arrows). Image brightness is saturated in the inset to reveal the chromatin bridge. (B) Nuclear cycle 1 (metaphase) with no apparent defect. (C) Nuclear cycle 2 with asynchronous nuclei and late segregating chromosome (arrow). (D) Nuclear cycle 2 (metaphase) with no apparent defect. (E) Aneuploid nuclear cycle 3 embryo. Note the aneuploid nuclear divisions with lagging chromosomes (arrows). (F) Nuclear cycle 3 (metaphase) with no apparent defect. (G-H) Confocal images of early *D. suzukii* embryos from a control cross 3 between infected Bal males and females. Normally developing embryos at (G) nuclear cycle 3 and (H) nuclear cycle 13. Maternally inherited Wolbachia cells are abundantly detected in the cytoplasm of these embryos. Chromosomes are labeled with anti-pan-histone (green), mitotic spindles are labeled with anti-alpha-tubulin (red) and *w*Suz *Wolbachia* are stained with DAPI (magenta). Note that the DAPI staining on Drosophila chromosomes was masked in the images (G, H) to facilitate visualization of bacteria (arrows). All scale bars: 20 µm.

### CI variation is explained neither by *Wolbachia* titer nor by the *Wolbachia* genome

Having established that *w*Suz induces CI, we next asked why its strength varied so markedly among lines, from undetectable (Rennes, Par6) to relatively strong (Bal). We first tested the bacterial density model, which predicts that CI strength increases with *Wolbachia* titer in testes (Breeuwer & Werren, 1993). Although *Wolbachia* titer in testes varied widely among the four *D. suzukii* lines (ANOVA: F₃,₁₆ = 6.17, P = 0.0055), with means ranging from 8 to 48.2 *Wolbachia* genome copies per host genome copy (Fig. 3), bacterial titer in testes did not explain variation in CI strength. Bal and Par6 did not differ in titer (ratio 0.96, Tukey-adjusted 95% confidence interval: 0.24-3.77; Tukey-adjusted P = 1.00; *SI Appendix*, Table S4) despite inducing 67% and no detectable CI, respectively. Conversely, Trento had the highest titer of all lines, about five-fold higher than Bal (ratio 5.61, Tukey-adjusted 95% CI: 1.42-22.07; P = 0.012; *SI Appendix,* Table S4), yet induced weaker CI (47%).

**Fig. 3.**
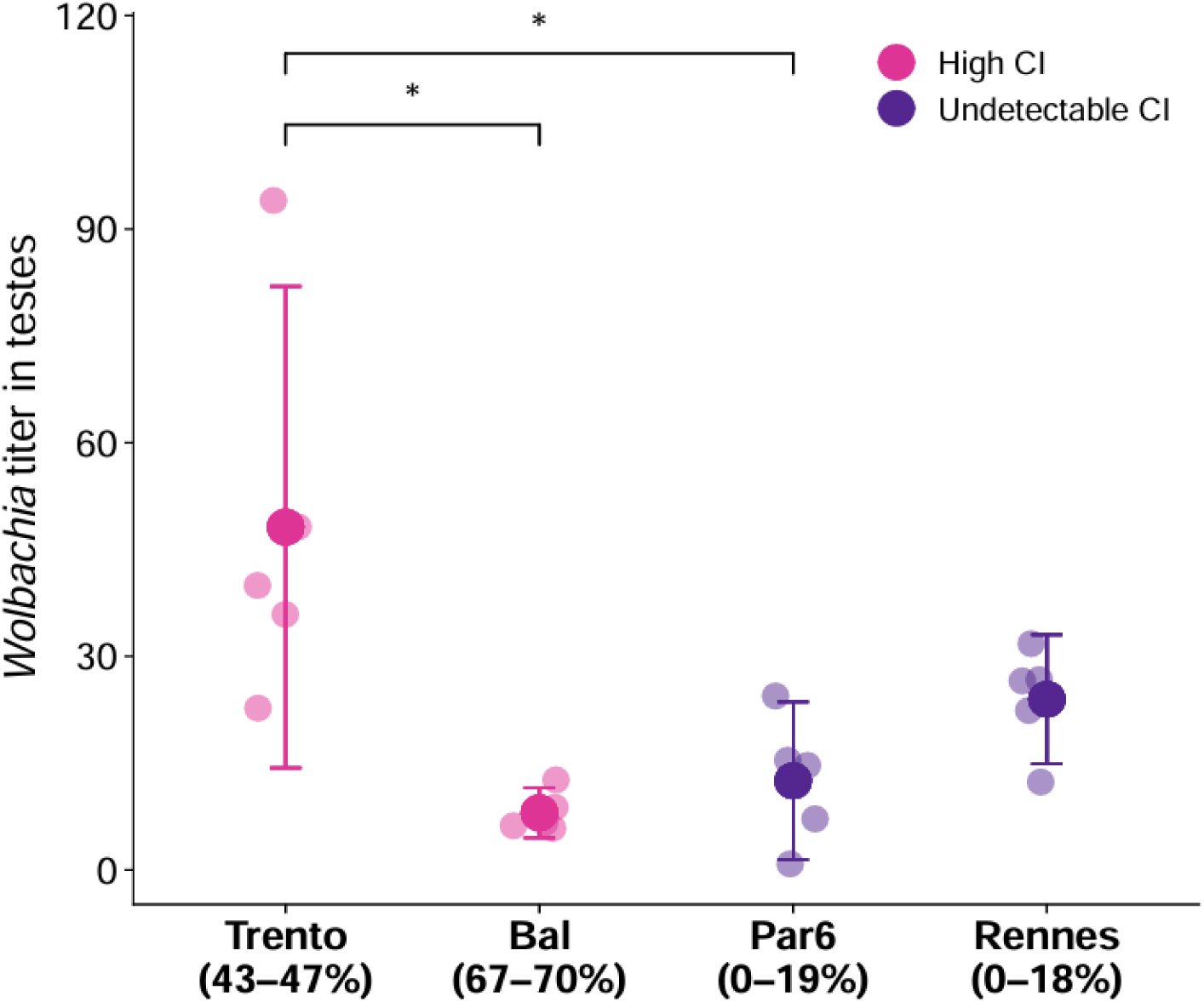
*Wolbachia* titer in testes of males from four *D. suzukii* lines. Titer in testes was calculated as 2^ΔCt^ of the *Drosophila* gene *RpL32* to the *Wolbachia* gene *ftsZ* and is expressed as the number of *Wolbachia* genome copies per host genome copy (n = 5 samples per line). Small circles show individual pools of male testes; large circles and bars show means and 95% confidence intervals. Color indicates whether males of each line induce CI. The strength of CI is given in parentheses below the line name. Significant differences are based on Tukey-adjusted pairwise contrasts on log_10_-transformed densities following ANOVA (F₃,₁₆ = 6.17, P = 0.0055); stars indicate significant comparisons (P < 0.05). Exact P values for all pairwise comparisons are reported in *SI Appendix* Table S4.

We next asked whether CI variation could instead stem from genetic differences among *Wolbachia* isolates, particularly in *cif* genes. Because the available *w*Suz reference genome, from Trento Province (Italy), is highly fragmented (Siozios et al., 2013), it does not allow a reliable characterization of multicopy regions such as *cif* loci. We therefore first assembled a new *w*Suz reference genome (wSuz_RW2025a; four contigs; *SI Appendix*, Fig. S2) from publicly available long-read whole-genome sequencing data of a *D. suzukii* sample from China (Wang et al., 2025). We found that this *w*Suz strain from China harbors three identical copies of Type I *cif* genes and a single copy of a Type II *cif* genes (Fig. 4A). Both Type I and Type II *cif* genes of *w*Suz showed low divergence from their homologs in related *Wolbachia* strains, but Type II *cif* genes clustered more closely with those of *w*Spc, which infects the sister species *D. subpulchrella*, than with those of *w*Ri from *D. simulans* (bootstrap support ≥ 67; Fig. 4B). Using this new reference genome, we next investigated whether the *Wolbachia* isolates differed genetically among the four lines of *D. suzukii*, particularly between the CI-inducing lines (Bal and Trento) and the non-CI-inducing lines (Rennes and Par6). Mapping short-read data from the four isolates onto the new assembly revealed very low genome-wide variation (Fig. 4C). All 18 sites fixed (allele frequency of 0 or 1) in each of the four isolates distinguished Rennes from the other three isolates, so none separated CI-inducing (Bal, Trento) from non-CI-inducing (Par6, Rennes) isolates. Moreover, all four isolates carried the same number of *cif* gene copies (Fig. 4D), with no SNP in any of them. All *w*Suz isolates carried one additional Type I *cifA–cifB* pair relative to *w*Ri (*SI Appendix*, Fig. S3).

**Fig. 4.**
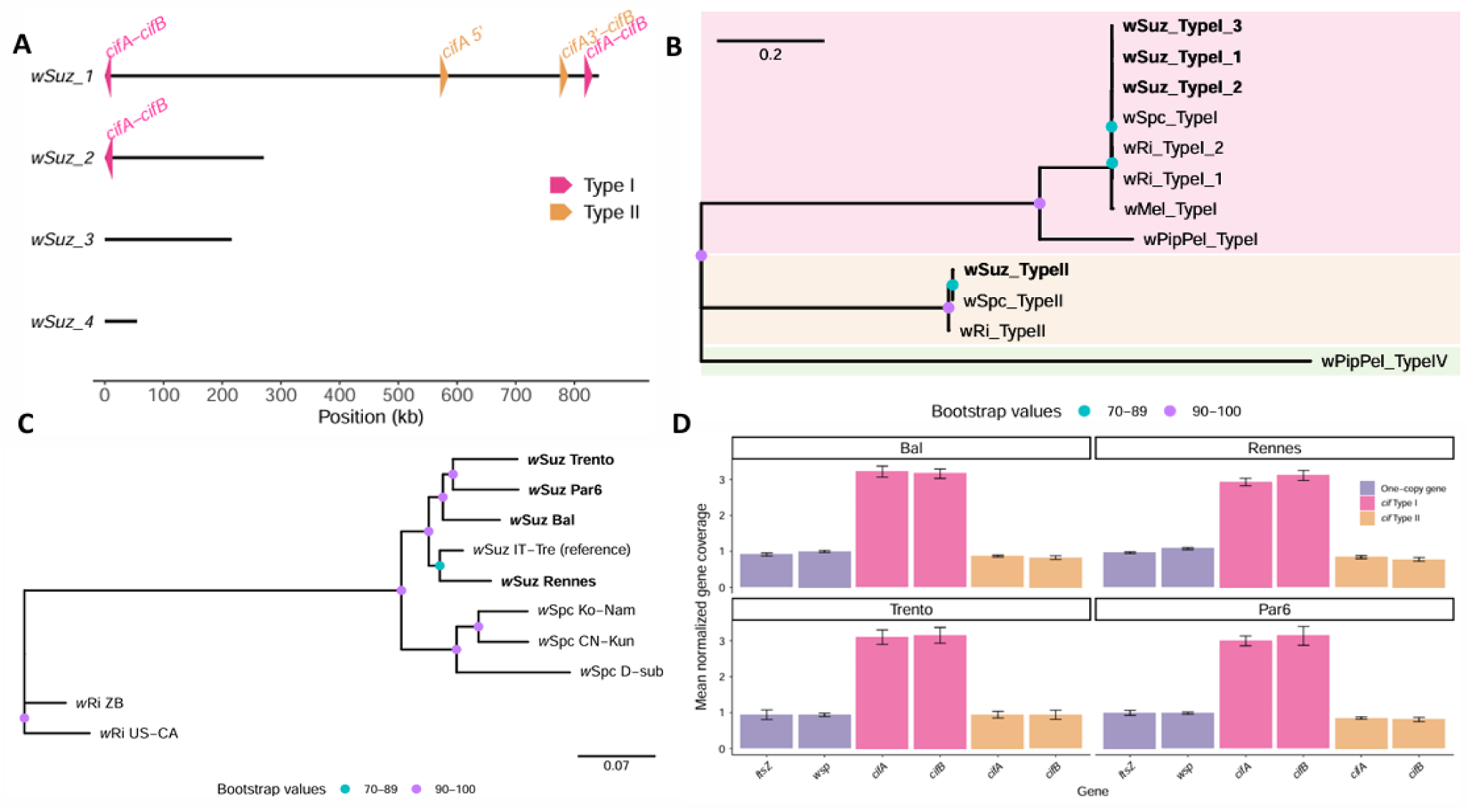
Genomic and phylogenetic characterization of *Wolbachia w*Suz and its *cif* genes. (A) Positions and types of *cif* loci identified along the four contigs (labeled *w*Suz_1–*w*Suz_4) of the new genome assembly of *w*Suz (*w*Suz_RW2025a). Type I and Type II *cif* loci are indicated by pink and orange symbols, respectively. The Type II *cifA* gene is split into two fragments by a ∼200 kb genomic rearrangement flanked by ISWpi7 insertion sequences. (B) Phylogenetic relationships among CifA–CifB sequences from *w*Suz and other *Wolbachia* strains from the sister species *D. subpulchrella* and the more distantly related *D. simulans* (*w*Ri), *D. melanogaster* (*w*Mel), *Culex pipiens* (*w*Pip). The phylogeny was inferred from 1704 amino acids (including 645 parsimony-informative sites) resulting from the concatenation of CifA and CifB sequences (JTT+F+G4 substitution model). Branch support values are indicated at the nodes. (C) Phylogenetic relationships among the ten *w*Ri-like genomes analyzed in this study, based on 554 SNPs. *Wolbachia* isolates sequenced in this study are shown in bold. Branch support values are indicated at the nodes. (D) Mean normalized sequencing coverage of *cifA* and *cifB* genes and two single-copy reference genes (*ftsZ* and *wsp*) in four *w*Suz isolates from *D. suzukii* lines that differ in the level of CI induced (Bal, Rennes, Trento, and Par6). Despite different levels of CI, *cif* genes of *w*Suz isolates had identical sequences to that of the reference *w*Suz_RW2025a assembly and harbored the same number of copies (3 copies of Type I *cif* genes and 1 copy of Type II *cif* genes). Error bars indicate SD in normalized sequencing coverage across gene positions.

### Additional crosses confirmed that the four *w*Suz isolates are functionally compatible

*w*Suz isolates were mutually compatible, and CI depends on the host nuclear background. Females carrying the Bal, Trento or Par6 *w*Suz isolate all rescued the CI induced by Bal or Trento males, consistent with the identical *cif* genes of these isolates. Thus, the Par6 isolate, in which CI was undetectable, remains fully functional for rescue (*SI Appendix*, Fig. S4, Table S5). Furthermore, introgressing the CI-inducing Bal and Trento isolates into the nuclear background of the uninfected Am line abolished CI, and *Wolbachia* titers became homogeneous among introgressed lines (*SI Appendix*, Figs. S5 and S6, Table S6). The Am nuclear background therefore suppresses CI, indicating that CI expression is controlled by the host nucleus rather than by the *Wolbachia* isolate.

### CI suppression is autosomal, partially dominant and involves few loci

We next investigated the genetic architecture of the observed host effect. We performed reciprocal crosses between the strong CI-inducing Bal line and the non-CI-inducing Par6 line, and performed CI assay for each paternal line, F1 sons and F2 grandsons. This design allowed us to test three alternative models to an additive autosomal model, which make distinct predictions for F1 and F2: dominant autosomal host suppressors, X-linked or Y-linked host suppressors (Fig. 5A–C). Hatch rates in test crosses with F1 males from both cross directions approached those with Par6 males, indicating that CI suppression is largely autosomal and involves both additive and dominant effects (Fig. 5D). The best-supported model included additive and dominance autosomal effects (full model, logit scale mean [95% confidence-interval]: additive effect, 1.02 [0.19, 1.94]; dominance effect, 0.57 [0.09, 1.12]; *SI Appendix*, Table S7b). Removing the additive or dominance effect from the best model decreased support (ΔAICc = 23.3 and 2.7, respectively; *SI Appendix*, Table S7a). The additive–dominance model accounted for 91% of the deviance explained by differences among cross types. The dominance effect was about 60% of the additive effect (*d*/*a* ≈ 0.56 on the logit scale; complete dominance corresponds to *d*/*a* = 1), consistent with incomplete dominance. We found no evidence for X- or Y-linked effects: adding either term to the best model did not improve its AICc, and both estimated effects were small, with 95% confidence intervals overlapping zero (full model, logit scale mean [95% confidence-interval]: X effect, 0.09 [−0.45, 0.62]; Y effect, −0.15 [−0.47, 0.16]; *SI Appendix*, Table S7b).

**Fig. 5.**
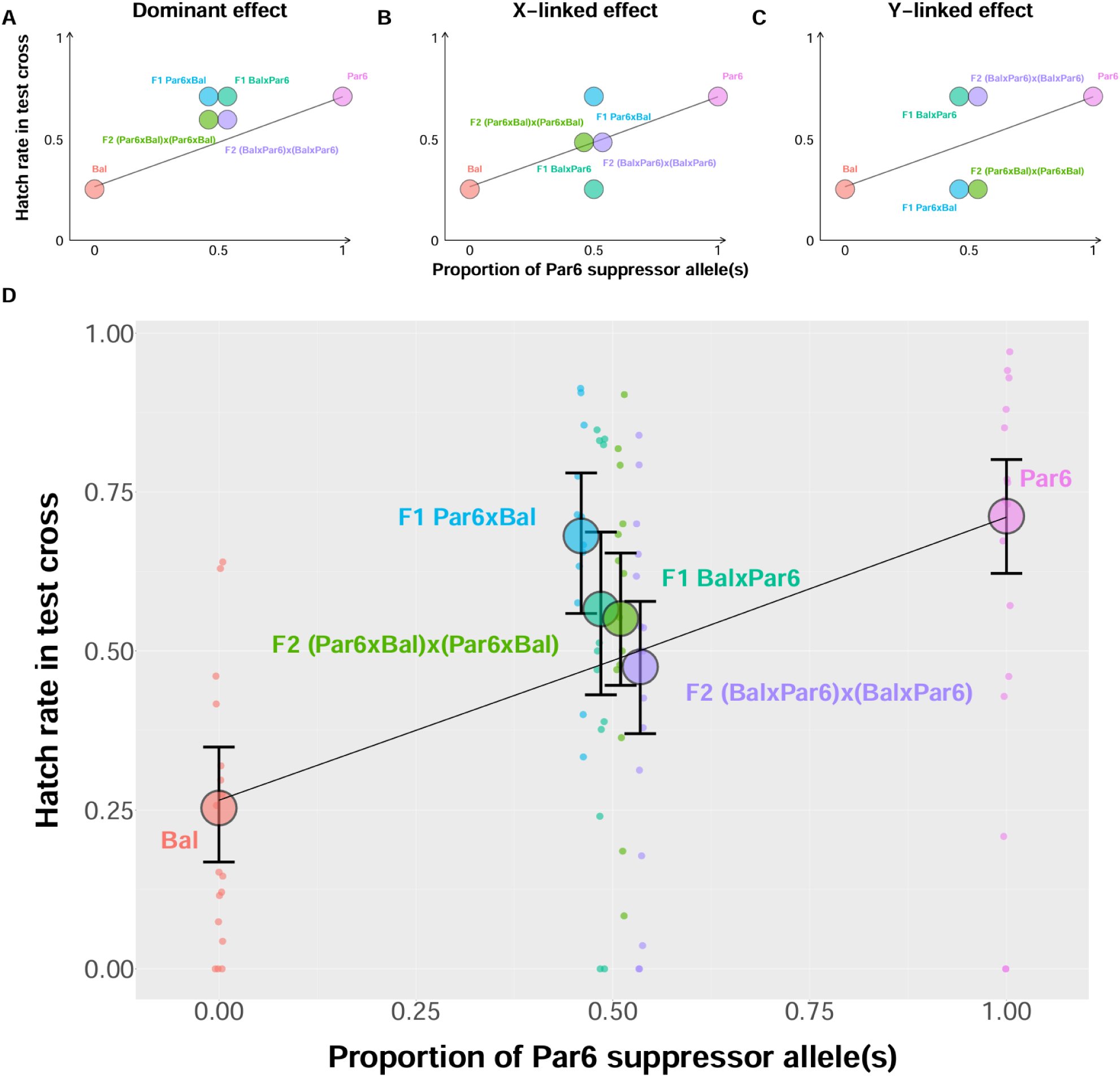
Expected (A-C) and observed (D) hatch rates in test crosses with parental, F1 and F2 males as a function of the proportion of host suppressor allele(s) derived from the Par6 parental line. Throughout the figure, crosses are denoted as female strain × male strain (e.g., Bal x Par6 indicates a Bal female crossed with a Par6 male). (A–C) Expected hatch rates in test cross under alternative models of CI suppressor genetic architecture (A) If Par6 suppressor allele(s) are autosomal with dominant genetic effects, F1 males are expected to exhibit the same level of sterility as males from Par6, whereas F2 males are expected to exhibit higher sterility due to increased homozygosity at non suppressor Bal allele(s) responsible for CI. (B) Under X-linked inheritance, F1 males are expected to exhibit the same level of sterility as males from their maternal lines, whereas F2 males are expected to show a level of sterility intermediate between those of the parental lines. (C) Under Y-linked inheritance, F1 and F2 males are expected to show the same hatch rate as males of the line from which their Y chromosome derives (paternal line for F1, paternal grandfather’s line for F2). (D) Observed hatch rates are consistent with autosomal suppressor allele(s) with partially dominant effects; X- and Y-linked effects were not detected. Fitted mean hatch rates and observed hatch rates for individual mating pairs are represented by large and small circles, respectively. Model selection was performed using AICc and 95%-confidence intervals, represented as error bars, were calculated using parametric bootstrap with 500 iterations (*SI Appendix*, Table S7). As a reference, diagonal solid lines represent the expected hatch rate of an autosomal model with additive genetic effects.

Hatch rates of F2 males spanned the full range between those of Bal and Par6 males (Fig. 5; *SI Appendix*, Fig. S7), and between-male variance was about fivefold higher among F2 males than among parental and F1 males. The Castle–Wright estimator (Castle, 1921) indicated a minimum effective number of 0.6 loci, i.e. consistent with a single locus (80% bootstrap interval: 0.4-2.7; 95% interval: 0.23-∞; 86% of estimates < 2 loci). The upper 95% bound was infinite because 2.6% of bootstrap samples showed no excess between-male variance among F2 males compared to parental and F1 males. Together, these results suggest that CI suppression is controlled by one or a few autosomal loci of major effect, with incomplete dominance of the Par6 allele(s), although a more polygenic architecture cannot be completely excluded.

### The genotype of uninfected females does not detectably modulate CI

We tested this in two ways. First, CI-inducing Bal males induced similar CI when crossed with uninfected females from three different lines (Am, Paris7 [Par7] or Montpellier47 [Mon47]; *SI Appendix*, Fig. S8). Second, non-CI-inducing Par6 and Rennes induced no detectable CI with uninfected Par7 and Mon47 females, respectively (*SI Appendix*, Fig. S8), so the absence of CI in these lines is not specific to the Am females used in our main assay. In both cases, allowing hatch rate to differ among female genotypes did not improve model fit (*SI Appendix*, Table S8).

## Discussion

*D. suzukii* has so far been considered a host in which *w*Suz fails to induce CI despite carrying *cif* genes (Cattel, Kaur, et al., 2016; Conner et al., 2017; Hamm et al., 2014). Our results resolve this apparent paradox. We found that its natural strain *w*Suz does induce CI, but that this phenotype is masked in host genotypes by partially dominant autosomal suppressors carried by males. CI variation ranged from undetectable to relatively strong among four European lines of *D. suzukii* and was explained by the nuclear genotype of males, but neither by variation in bacterial titer nor by variation in the *w*Suz genome, which was virtually identical among isolates. These findings shift the explanation for weak or absent CI from the symbiont to the host, and suggest that host suppression may be a widespread yet overlooked determinant of variation in CI expression in nature.

### Neither *Wolbachia* titer in male testes nor *Wolbachia* genotype explained the variation in CI strength among host lines

According to the bacterial density model, higher bacterial titers are expected to result in stronger CI (Breeuwer & Werren, 1993). However, we found no effect of *Wolbachia* titers in testes on CI strength, which is in line with results in other *Drosophila-Wolbachia* systems (Bagchi et al., 2026; Martinez et al., 2015; Shropshire et al., 2022; Yamada et al., 2007). A second potential source of variation in CI is the *Wolbachia* genotype itself. Variation in *Wolbachia* genotype could potentially contribute to CI differences through variation in the presence, copy number, or sequence of *cif* genes (Bonneau et al., 2018; Martinez et al., 2021). Our genomic analyses provided no evidence that CI variation is driven by *cif* copy number or sequence polymorphism. All four *w*Suz isolates were identical to the new reference genome at the three Type I copies and the single Type II copy. Short reads may miss structural variants near *cif* loci that could alter expression, but this seems unlikely because the same isolates lose CI in the Am nuclear background.

### Male, not female, host genotype governs CI strength

Strong variation in *Wolbachia*-induced CI among host genotypes has been documented when a *Wolbachia* strain is transferred between host species (Bordenstein et al., 2003; Boyle et al., 1993; Martinez et al., 2015; Poinsot et al., 1998), and there is growing evidence that variation of similar magnitude occurs within species. Mean CI ranges from 30% to 88% among *D. melanogaster* genotypes (Reynolds & Hoffmann, 2002), from 16% to 100% among *Tetranychus urticae* genotypes (Wybouw et al., 2022) and from about 25% to 100% between *Callosobruchus analis* genotypes (Numajiri et al., 2025). Our results extend this pattern to *D. suzukii*, where CI ranged from undetectable to 67% across the four host lines tested, a range comparable to those reported in *D. melanogaster* and *T. urticae*. Nikolouli et al., 2020 suggested that host genotype also modulates CI in *D. suzukii* lines transinfected with *w*Tei, although CI remained strong in all of them (64-96%). *w*Tei carries a truncated Type I *cifB* but an apparently intact Type IV *cifA-cifB* pair (Baião et al., 2021). Whether *D. suzukii* suppressors may have different efficacy against the Type I CifB of its native strain *w*Suz and the Type IV CifB of *w*Tei remains to be investigated.

Beyond its magnitude, the source of this variation is noteworthy: CI strength depended on the genotype of infected males, whereas the genotype of uninfected females had no detectable effect. This suggests that host-driven variation in CI arises predominantly from processes operating in the male germline during toxin production or during its loading into maturing sperm nuclei prior to fertilization (Horard et al., 2022), rather than from maternal factors in the egg. Few studies have compared how male and female host genotypes contribute to CI. In *D. simulans*, CI shows no detectable host genetic variation (Carrington et al., 2011; Hoffmann & Turelli, 1988), whereas it varies with the nuclear background of infected males in *D. melanogaster* and *D. teissieri* (Cooper et al., 2017; Reynolds & Hoffmann, 2002). In the spider mite *Tetranychus urticae* and the bean beetle *Callosobruchus analis*, both sexes contribute to CI variation, but male genotype has the stronger effect (Numajiri et al., 2025; Wybouw et al., 2022).

Understanding differences in the relative contribution of male and female genotypes first requires knowing how host suppression is genetically encoded: how many loci are involved, where they are located in the genome, and whether their effects are dominant or recessive. Yet the genetic architecture of host suppressors of CI remains poorly characterized (but see Cooper et al., 2017; Numajiri et al., 2025; Wybouw et al., 2022). We provide a first answer for *D. suzukii*. Our crosses indicate that CI suppression in *D. suzukii* is partially dominant and mainly autosomal: F1 sons induced little CI, and CI partially reappeared in F2 sons. Castle–Wright estimates suggest that this suppression involves a small number of loci (minimum number of effective factors ≈ 0.6; 80% confidence interval: 0.4-2.7 loci), comparable to the two to five loci estimated for the bean beetle *C. analis* (Numajiri et al., 2025) and consistent with the polygenic basis of CI suppression in the spider mite *T. urticae* (Wybouw et al., 2022). Because the Castle–Wright estimator assumes unlinked loci of equal, additive effect, it gives only a lower bound, and genetic mapping will be needed to determine the exact number and location of suppressor loci.

Host suppressors may act through different mechanisms, which include the evolution of host factors that suppress *Wolbachia* or its CI-inducing toxin or divergences of the host pathways targeted by the toxin (Shropshire et al., 2020). Although host genetic suppressors of *Wolbachia* density, and consequently toxin production, are obvious candidates, our results provide no support for such a mechanism. Suppression did not act through bacterial density and did not involve the female genotype. It therefore most likely operates in the male germline, downstream of *Wolbachia* proliferation: by reducing *cif* expression, by limiting the loading of CifB into maturing sperm nuclei (Horard et al., 2022), or by neutralizing the toxin before fertilization. Investigating these possibilities defines the next step in understanding how hosts evolve to control the toxin delivered by their symbionts.

### Evolutionary dynamics of host suppressors of CI

Theoretical models predict that host suppressors of CI spread because infected carrier males escape CI when mating with uninfected females; their spread is therefore driven by the frequency of such crosses (Kauai & Wybouw, 2025; Koehncke et al., 2009; Turelli, 1994). Selection on suppressors is expected to weaken, but not vanish, as *Wolbachia* approaches fixation, because imperfect maternal transmission keeps producing uninfected females; under these conditions suppressors spread only over hundreds of generations (Kauai & Wybouw, 2025). Selection should be stronger in species such as *D. suzukii*, where infection frequencies remain intermediate and variable among populations (Cattel, Kaur, et al., 2016). In turn, by weakening CI-driven spread, suppressors may help maintain these intermediate frequencies. Across the different genotypes of *D. suzukii* assayed in this and previous studies (Cattel, Kaur, et al., 2016; Hamm et al., 2014), most genotypes showed no CI and even the uninfected Am line carried suppressors (SI Appendix, Fig. S6), which suggests that suppressor alleles may occur at high frequencies in natural populations. Identifying suppressor alleles would make it possible to track their spatial and temporal dynamics in natural populations of *D. suzukii* and to test whether their frequency correlates with that of *Wolbachia*. The same question arises at a broader scale: why do some host species harbor suppressors while others do not? Host suppressors of CI appear widespread among insect and mite hosts (Fig. 6), yet a few species, such as *Culex pipiens* and *D. simulans*, show no evidence of host suppression (Carrington et al., 2011; Hoffmann et al., 1986; Sicard et al., 2021). To reconcile these contrasting observations, we propose a threshold model in which the level of cytoplasmic incompatibility is determined by the interaction between the *Wolbachia* genotype and the host genotype. In this model, differences among host genotypes in their ability to suppress CI remain phenotypically invisible in two situations: when *Wolbachia* produces no toxin, so that no host can express CI (e.g., *w*Au, which lacks functional *cif* genes, in *D. simulans*), and when *Wolbachia* produces a very toxic CifB so that CI remains complete even in hosts that might harbor suppressor alleles (e.g., potentially *w*Pip in *Culex pipiens*; Fig. 6). For some strains of *Wolbachia*, differences among host genotypes result in variation in the quantity of toxin that spans the threshold required to induce embryo mortality (Fig. 6). For a given combination of host and *Wolbachia* genotypes, the quantity of toxin can additionally vary among males due to environmental factors (e.g. temperature, male development time, male age, etc.; Bagchi et al., 2026; Reynolds & Hoffmann, 2002; Shropshire et al., 2021; Yamada et al., 2007).

**Fig. 6.**
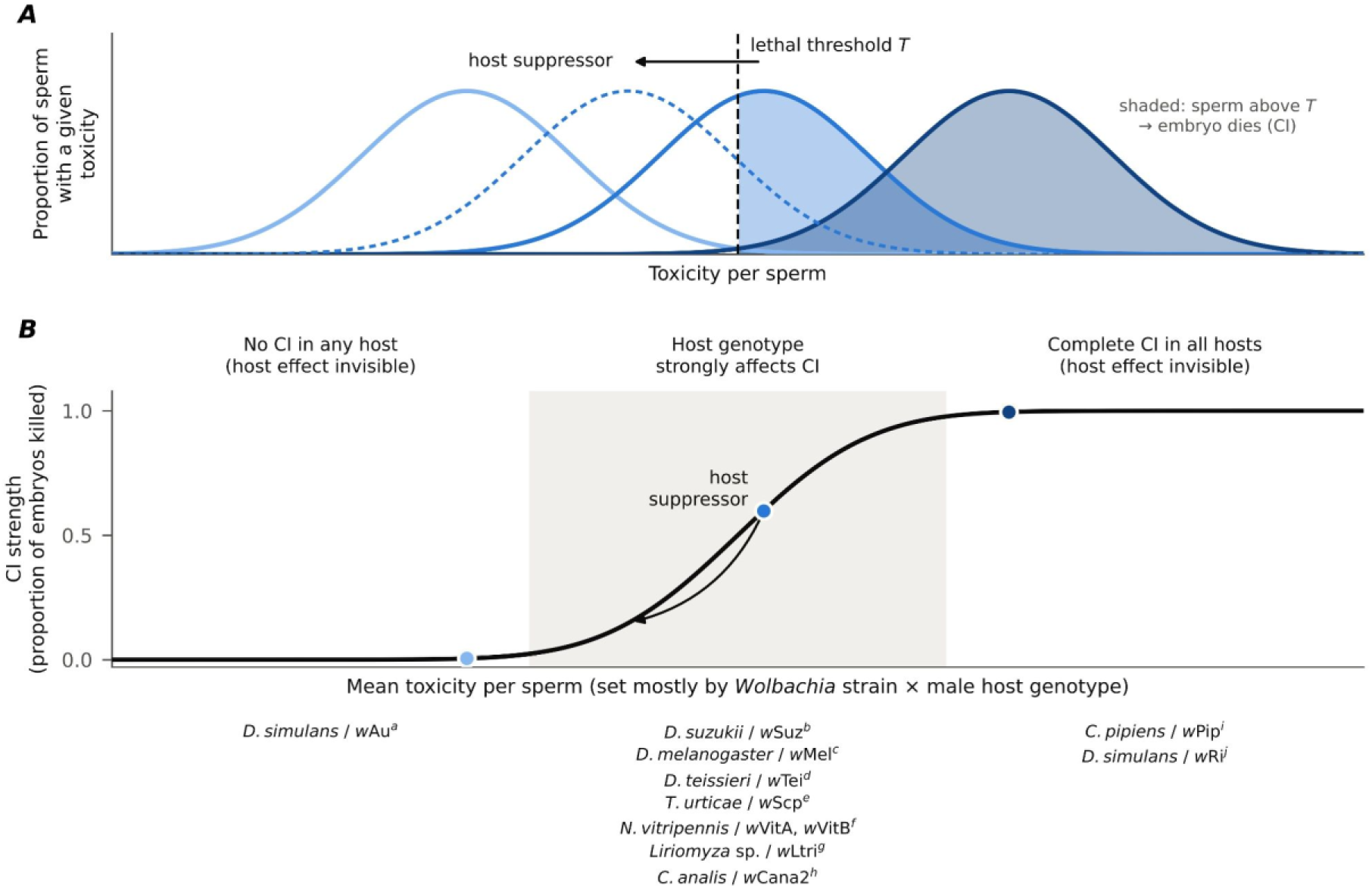
A threshold model for how host genotype affects CI strength. (A) Toxicity, the effectiveness of the toxin, varies among sperm of infected males around a mean set jointly by the *Wolbachia* strain and the male host genotype. An embryo dies when toxicity exceeds a threshold *T* (dashed line), so CI strength equals the proportion of the distribution above *T* (shaded). Host suppressors shift the distribution to the left. Environmental factors, such as male age or temperature can also shift the distribution to the right or to the left. (*B*) CI strength as a function of the mean toxicity. Because this relationship is sigmoid, a given shift caused by host suppressors strongly changes CI only when the mean lies near the threshold (central zone). It has little effect when the mean is far below the threshold (no CI in any host) or far above it (complete CI in all hosts). Host–*Wolbachia* associations are placed according to their observed CI and host effects, not measured toxicity: (a) Hoffmann et al., 1996, (b) This study, (c) Reynolds & Hoffmann, 2002, (d) Cooper et al., 2017, (e) Wybouw et al., 2022, (f) Bordenstein & Werren, 1998, (g) Ohata et al., 2025, (h) Numajiri et al., 2025, (i) Sicard et al., 2021, (j) Hoffmann et al., 1986. The strong CI of *w*Pip and *w*Ri may reflect high toxicity that exceeds the threshold in all host genotypes, or the absence of suppressor alleles in their host populations.

### Implications for *Wolbachia*-based control of *D. suzukii* and other pests and vectors

*D. suzukii* is one of the most damaging invasive pests of soft and stone fruits worldwide, and the limitations of insecticide-based control have prompted the development of *Wolbachia*-based alternatives. The Incompatible Insect Technique (IIT) consists of releasing *Wolbachia*-infected males whose matings with wild females yield inviable embryos through CI. Combined with irradiation, it has successfully suppressed mosquito populations in the field (Bansal et al., 2024). It has also been under development for *D. suzukii* using the transinfected strain *w*Tei, which is incompatible with both uninfected and *w*Suz-infected females (Nikolouli et al., 2020). Our results bear on these strategies in three ways.

First, because male genotype modulates CI strength, the efficacy of IIT depends on the genetic background of the released males and not only on the *Wolbachia* strain they carry. Release strains should therefore be screened for suppressors, or purged of them by artificial selection. Because suppression is partially dominant, a single suppressor allele introduced when mass-reared strains are refreshed with wild individuals – a common practice to limit inbreeding – could be enough to erode CI, so CI strength should be monitored over time. Notably, *w*Tei induced strong CI in two *D. suzukii* genotypes tested (Nikolouli et al., 2020), which suggests that it may be less sensitive to these suppressors than *w*Suz. Second, we detected no effect of female genotype, although with three female lines our power is limited. If confirmed, this would suggest that variation among wild females offers little scope for resistance to CI, although female effects on CI have been reported in spider mites and beetles (Numajiri et al., 2025; Wybouw et al., 2022).

Third, host suppressors threaten strategies that use CI to spread *Wolbachia* through wild populations, such as population replacement (Hoffmann et al., 2011). Theory predicts that suppressors are favored mainly when infected males frequently mate with uninfected females, that is, when *Wolbachia* is at intermediate frequency (Koehncke et al., 2009; Turelli, 1994). Replacement programs necessarily pass through this phase, and suppressors selected during it could slow the spread of *Wolbachia* or prevent it from reaching fixation. Once *Wolbachia* is near fixation, selection on suppressors weakens but does not vanish, because imperfect maternal transmission continues to produce uninfected females. Suppressor alleles could therefore persist and weaken the CI needed to offset that imperfect transmission. Mapping suppressor loci would provide markers to select release strains and to monitor suppressor frequencies in target populations.

### Conclusion and perspective

Our study provides strong evidence that naturally occurring male host genetic suppressors are major determinants of variation in *Wolbachia*-induced CI in *D. suzukii*. Together with previous studies that failed to detect CI, our results suggest that host suppressors are widespread in natural populations and represent an important, yet largely overlooked, component of the evolutionary dynamics of CI. Future studies should identify the genes and molecular mechanisms responsible for suppressing CI and the dynamics of suppressor alleles across natural populations. Beyond their evolutionary significance, these findings also have practical implications for the development of new control strategies for insect pests or vectors based on *Wolbachia*.

## Materials and Methods

### Fly rearing and cytoplasmic incompatibility (CI) assay

PCR-based screening of 18 *D. suzukii* lines collected across European sites between 2011 and 2018 and maintained in the laboratory since, identified four *Wolbachia*-infected *lines* (Bal, Trento, Rennes and Par6; *SI Appendix*, Table S1). All lines were maintained on “sick fly” medium (Backhaus et al., 1984) at 17±2°C with 60±2% relative humidity (16:8 light:dark cycle). Prior to conducting CI assays, lines were acclimated in a climate-controlled chamber at 21±2°C and 60±2% relative humidity (16:8 light:dark cycle) for at least two generations (CI assays were performed under the same conditions).

For CI assays, we compared hatch rate between test crosses and three different types of control crosses (to control for potential confounding effects such as heterosis, maternal or paternal genetic effects) using single-pair mating following Hamm et al., 2014 (Fig. 1A). Ten to twenty virgin females were collected 4-6 hours post-emergence in tubes with medium and allowed to mature for five days. Before each cross, the virgin status was confirmed by checking tubes for the absence of larvae. For each cross, a single pair of 5-day-old virgin female and 2-3-day-old male from infected and/or uninfected *D. suzukii* lines were placed together in a small embryo-collection cage (Genesee Scientific, USA) with a Petri dish filled with 4 mL of blackberry medium. This medium was made using a recipe modified from Olazcuaga et al., 2019 (1092 g of blackberry purée; 816 mL of 2.5% agar mixed in water; 108 g of malted yeast; 27 g of yeast extract; 18 mL of ethanol; 1.8 g of nipagin; and 10.8 mL propionic acid). Because males were 2–3 days old at mating, we tested whether male age within this range affected hatch rate in test and compatible crosses (*SI Appendix*, Table S9a-c). For each type of cross, we performed 13-76 independent single-pair matings. Each pair of flies was transferred daily onto a Petri dish with fresh medium for 3 or 5 days. After each transfer, we assessed the fecundity of each female by counting the number of eggs laid over 24 hours. To allow embryos to hatch and larvae to develop, Petri dishes were incubated for 48 additional hours. We counted the number of larvae in each Petri dish after extracting them by adding 1 mL of 20% sucrose solution (a.k.a. sugar flotation technique; Shaw et al., 2019). We did not distinguish between unfertilized eggs and embryos that died during development (Hamm et al., 2014).

All statistical analyses of CI assays were performed using the R Statistical Software (v4.3.2, R Core Team, 2023). We excluded pairs whose females laid no eggs from the analyses. When the number of larvae was greater than the number of eggs counted, we set the number of eggs to the number of larvae counted. For each pair, hatch rate was calculated as the ratio of larvae to the total number of eggs laid.

As previously observed in Hamm et al., 2014 in *D. suzukii*, a substantial proportion of mating pairs produced eggs but no larvae. These zero-hatch cases also occurred in control crosses between uninfected flies and therefore could not be attributed solely to CI. To account for this excess of zeros, which is likely due to female sterility or failed mating, we compared hatch rates between test and control crosses using zero-inflated beta-binomial models (R package glmmTMB; Brooks et al., 2017). For each infected line, the response was the proportion of eggs that hatched per mating pair, weighted by the number of eggs laid. The conditional (beta-binomial) part of the model tests for differences in hatch rate among cross types. We fitted three models (*SI Appendix*, Table S2). (i) In the null model, the mean hatch rate was constant across the four types of crosses. (ii) In the CI model, the mean hatch rate differed only between the test cross (uninfected females × infected males) and the three control crosses, which shared a common mean. (iii) In the saturated model, the mean hatch rate differed among the four types of crosses. All models included a constant zero-inflation parameter (accounting for variation among pairs in the probability of producing no larvae) and a constant dispersion parameter (accounting for overdispersion in hatch rate among mating pairs). Because the saturated model can be best supported simply because the three control crosses differ in hatch rate (e.g., through heterosis), it does not by itself indicate CI. We therefore considered that a line induced CI when either (i) the CI model was best supported, or (ii) the saturated model was best supported and the 95% confidence intervals of all three CI estimates, each calculated with a different control cross as the reference (see below), excluded zero.

To select for the best model, we used the corrected Akaike Information Criterion (AICc; Burnham & Anderson, 2002). Models with AICc differences < 2 compared to the model with the lowest AICc are considered as equally supported by the data (ΔAICc < 2). We computed the 95% confidence interval of each type of cross using parametric bootstrap with 500 iterations (Table S2; percentile bootstrap method; Davison & Hinkley, 1997). In addition, we also computed three estimates of CI using each of the three control crosses as a reference using the formula of Poinsot et al., 1998. Finally, to ensure that our egg and larvae counts were accurate, we computed repeatability of the log-transformed embryo and larvae counts (Zriki et al., 2024). Repeatability analyses of egg and larval counts were high in our experiments, with intra-observer repeatability of 99% and 96%, respectively, and inter-observer repeatability of 99% and 93%, respectively.

### Test for cytoplasmic incompatibility in the four lines of *D. suzukii*

To test for cytoplasmic incompatibility in the four lines of *D. suzukii* infected with *Wolbachia* (Bal, Trento, Rennes and Par6), we used the CI assay described above. The experiment was split over 2023 and 2024 with ten temporal blocks of one week each, with replication of crosses among some of the blocks. To test whether the reduction in hatch rate observed in test crosses with Bal and Trento males (see results) was due to *Wolbachia* or another bacterium, we reared these two lines and the line from Rennes (as a control) on “sick fly” medium supplemented with 0.25 mg/mL of the antibiotic tetracycline for three generations (Fry et al., 2004). Following antibiotic treatment, the antibiotic-treated lines were tested by PCR, and *Wolbachia* was not detected. We then performed a CI assay (*SI Appendix,* Fig. S1).

### Cytological analysis of early embryos

For cytological analyses, 100 embryos from the test (Am females × Bal males) and control cross 3 (Bal females × Bal males) were collected on agar plates with forceps and transferred in a small basket and dechorionated in 20% of bleach. After rinsing in water, they were briefly dried and then placed in a 1:1 mixture of heptane:methanol at −20°C. Fixed embryos were rinsed 3 times in methanol and rehydrated in PBS 1X, 0.15% Triton X-100, (PBST) 3 × 20 minutes. Embryos were then incubated with primary antibodies in PBST, overnight at +4°C on a rotating wheel. After washing three times in PBST, embryos were incubated with secondary antibodies following the same procedure. After the final wash, embryos were mounted in mounting medium (DAKO, ref #S3023) containing 1μg/mL DAPI. Primary antibodies: mouse anti-histone IgG2a antibody (1:1000; Millipore MABE71), mouse anti-alpha tubulin IgG1 clone DM1A (1:500, Sigma Aldrich T9026). Secondary antibodies: AlexaFluor 488 anti-mouse IgG2a (1:1000, Invitrogen A21131) and AlexaFluor 555 anti-mouse IgG1 (1:1000, A21127). Images were acquired on an LSM800 confocal microscope (CarlZeiss) equipped with a X40 (1.4 N.A.) objective lens and processed using the Zen (CarlZeiss) and Fiji softwares.

### Effect of *Wolbachia* titer on cytoplasmic incompatibility

To test whether CI strength is associated with *Wolbachia* titer, we quantified *Wolbachia* genome copies relative to host genome copies by TaqMan qPCR in the testes of 3-day-old males from each infected line (five pools of five testis pairs per line). We used *ftsZ* for *Wolbachia* and the single-copy *Rpl32* gene for the host (Cattel, Martinez, et al., 2016). We calculated titer as 2^ΔCt^ (Hague et al., 2020), *where* ΔC_t_ = C_t*Host*_− C_t*Wolb*_ *, C_tWolb_* and *C_tHost_* represent cycle threshold values for the *Wolbachia ftsZ* and Drosophila *Rpl32* amplification, respectively. *Wolbachia* titers were log_10_–transformed to meet the assumptions of normality and homoscedasticity, verified by inspection of residual and quantile–quantile plots, and compared among lines by ANOVA followed by Tukey-adjusted pairwise contrasts on the log scale (R package emmeans; Lenth, 2024). Details are given in *SI Appendix*.

### New draft genome assembly of *w*Suz and phylogeny of *cif* genes

We assembled a new *w*Suz reference genome (Zenodo, https://doi.org/10.5281/zenodo.24090511) from publicly available PacBio long-read data (SRA SRX27255168; Wang et al., 2025; *SI Appendix*; Fig. S2). We retrieved *cif* genes by BLAST searches with *w*Ri *cif* sequences. We then inferred a maximum-likelihood tree with IQ-TREE 3 (Wong et al., 2026) from concatenated, aligned CifA and CifB protein sequences of *w*Suz, *w*Sp*, w*Ri, *w*Mel and *w*Pip (Martinez et al., 2021; *SI Appendix*).

### Genetic diversity among *Wolbachia* isolates from the four lines of *D. suzukii*

To genotype the four *w*Suz isolates and assess *cif* polymorphism, we extracted DNA from the ovaries of 25 females per line, enriched for endosymbionts by sequential filtration (*SI Appendix*). We then sequenced it on an Illumina NovaSeq 6000 (2 × 250 bp; ENA PRJEB127904). We added published reads from *w*Suz (Trento; Siozios et al., 2013), *w*Spc and *w*Ri (*SI Appendix*, Table S10). After filtering with fastp (Chen et al., 2018), we mapped reads to the *w*Suz reference genome together with the *D. suzukii* nuclear and mitochondrial genomes (Camus et al., 2025). We called variants jointly with bwa-mem2, SAMtools and FreeBayes. We computed pairwise divergence among the ten *Wolbachia* genomes with poolfstat (Gautier et al., 2022), keeping positions present in at least 90% of genomes. From this divergence, we inferred a neighbor-joining tree with ape (Paradis & Schliep, 2019), with 1,000 bootstrap replicates. We estimated *cif* copy number as each gene’s mean read depth relative to the mean depth of its contigs using SAMtools (Li et al., 2009).

### Testing for functional differences among *Wolbachia* isolates using rescue assays and introgressions of *Wolbachia* isolates

To test whether the identical *cif* genes of the different *w*Suz isolates allow mutual rescue, we crossed Bal with each of two other infected lines, Trento and Par6 (*SI Appendix*, Fig. S4). To test for a cytoplasmic effect on CI, we introgressed the cytoplasm (including *Wolbachia*) of the four lines whose males induce different levels of CI (Bal, Trento, Rennes and Par6) into the nuclear background of the uninfected Am line, using seven generations of backcrossing (following Cooper et al., 2017). The four resulting lines (Am^Par6^, Am^Rennes^, Am^Trento^, Am^Bal^) carry different cytoplasms but almost identical nuclear backgrounds, each more than 99% Am background. We measured *Wolbachia* titer and CI in these introgressed lines (*SI Appendix*, Figs. S5 and S6).

### Relative roles of *D. suzukii* male and female genotypes in cytoplasmic incompatibility

To quantify the contribution of male genotype to *Wolbachia*-induced CI, we performed reciprocal crosses between the two *Drosophila suzukii* lines with the strongest (Bal) and weakest (Par6) CI phenotypes. We then measured CI induced by males from the parental lines and from the F1 and F2 generations, each mated with uninfected Am females. Each couple contributed two to three Petri dishes (229 dishes from 87 couples).

Hatch rates were analysed with zero-inflated beta-binomial generalized linear mixed models with a constant zero-inflation probability, with couple as a random intercept to account for repeated dishes per couple (*SI Appendix*). Mean hatch rates for each cross type were estimated from a model with cross type as a fixed effect. To test whether differences in hatch rate between cross types reflected additive autosomal, dominant autosomal, X-linked or Y-linked effects, we used a generation means approach (Gilchrist & Partridge, 1999). We fitted 15 models combining these four genetic parameters, including an intercept-only model (*SI Appendix*, Table S7). Models were ranked using the Akaike information criterion corrected for small sample size (AICc; Hurvich & Tsai, 1989).

To further determine whether female genotype could also impact CI, we compared the hatch rates in crosses either between Bal males (which induce CI, see results) and females from three uninfected lines (Am, Mon47, Par7) or between Par6 and Rennes males (which do not induce CI, see results) and uninfected females from different lines (*SI Appendix*, Table S8).

## Data availability

All sequencing reads were deposited in the European Nucleotide Archive (ENA; project PRJEB127904). The *w*Suz_RW2025a whole genome assembly is publicly available on Zenodo (https://doi.org/10.5281/zenodo.24090511). Phenotypic data and analysis scripts are available at https://github.com/luchiksveta05/Y2025.Wolbachia.CI.

## Acknowledgments

We wish to thank Bregje Wertheim, Patricia Gibert, Laurence Mouton and Marc Vreysen for providing some of the strains of *D. suzukii*; Hélène Henry for providing qPCR probes; Luis Teixeira, Ewa Chrostek, Nicky Wybouw, Emmanuelle Jousselin and Anne Laure Clamens for discussions and Stéphanie Chalmeton and Florian Cros from the Drosophila Biocampus facility for their help with medium preparation. We are also grateful to the Genotoul bioinformatics platform Toulouse Occitanie (Bioinfo Genotoul, https://doi.org/10.15454/1.5572369328961167E12) for providing computing resources and the Lyon SFR Biosciences (UAR3444/CNRS, US8/INSERM, ENS de Lyon, UCBL) imaging facility (PLATIM). Svitlana Serga was supported by the PAUSE-ANR Ukraine Program. Mountassira Matin and Chooi Lin Phooi were supported financially by the IDIL Graduate Program of the University of Montpellier funded by the French Government (France 2030) via the Agence Nationale de la Recherche (ANR-21-SFRI-0004). This project has received funding through the Agence Nationale de la Recherche (ANR-22-CE20-0002-01) and the MSCA4Ukraine project funded by the European Union. Views and opinions expressed are, however, those of the author(s) only and do not necessarily reflect those of the European Union. Neither the European Union nor the MSCA4Ukraine Consortium as a whole nor any individual member institutions of the MSCA4Ukraine Consortium can be held responsible for them.

## Supporting Information

### Appendix S1 *Wolbachia* detection in the different *D. suzukii* lines

We screened 18 available lines of *D. suzukii* for *Wolbachia* infection, including those established from European populations (Italy and France; Table S1). DNA was extracted from a pool of five random live or ethanol-preserved females from each line using the DNeasy Blood & Tissue Kit (Qiagen, USA) according to the manufacturer’s recommendations. To detect *Wolbachia* infection, we performed standard multiplex PCR using primers targeting the *Wolbachia surface protein* gene (*wsp* primers 81F/691R; Zhou et al., 1998), the bacteria-specific *filamenting temperature sensitive* gene (*ftsZ* primers ftsZ-F2/ftsZ-R2; Cattel et al., 2016), and a host microsatellite (DS-14, primers; Fraimout et al., 2015).

The PCR reaction mix was prepared using the Multiplex PCR Kit (Qiagen, USA) in a total volume of 10 µL containing 5 µL of QIAGEN Multiplex Master Mix, 0.2 µL of 10 µM for each primer (to get a final concentration of 0.2 µM), and 2 µL of the DNA template for each strain tested. Cycling conditions start at 95°C (15 mins) to activate the HotStart Taq DNA polymerase, followed by denaturation at 94°C (30s), 57°C (90s) for annealing, extension at 72°C (90s) for 35 cycles, with a final extension at 72°C (10 mins). PCR products were visualized using 1.5% agarose gel electrophoresis. To ensure the absence of DNA contamination, we included a negative control, consisting of a reaction with water instead of DNA. A positive control was the line Od107 of *D. simulans* known to be infected with *Wolbachia* (Serga et al., 2023).

### Appendix S2 Males from antibiotic-cured lines do not induce cytoplasmic incompatibility

To test whether the reduced hatch rate in the test crosses was caused by *Wolbachia*, we removed *Wolbachia* (and other bacteria) from the Bal, Trento and Rennes lines by rearing them on German food medium with tetracycline (0.25 mg/mL) for three generations (Fry et al., 2004). To limit direct effects of the antibiotic, the treated lines (hereafter Bal^T^, Trento^T^, and Rennes^T^) were then maintained on standard medium for three generations before the experiments. The effectiveness of the antibiotic treatment was confirmed by PCR; all treated lines were negative for *Wolbachia*. Hatch rates were then measured in test and control crosses in a separate block of experiments (block 5).

We analysed hatch rates with zero-inflated beta-binomial models as described in the Materials and Methods (Table S3a). In Bal^T^, the null model, with no difference among cross types, ranked first (Table S3a, c), indicating an absence of CI. In Trento^T^, the model contrasting the test cross with the control crosses ranked first, but it was less than two AICc units better than the null model (ΔAICc = 1.78), which does not constitute evidence for a reduced hatch rate in the test cross. In Rennes^T^, the saturated model ranked first (ΔAICc = 1.90 relative to the null model). This reflected differences among control crosses in this line: the hatch rate of the test cross (mean estimate: 0.768, 95% confidence interval 0.682–0.837) did not differ from that of the uninfected control cross (mean estimate: 0.756, 95% confidence interval0.673–0.831) (Table S3b, Fig. S1). These results indicate that the reduction in hatch rate in the test cross of naturally infected lines depends on a bacterial agent removed by tetracycline, consistent with *Wolbachia*.

**Fig. S1.**
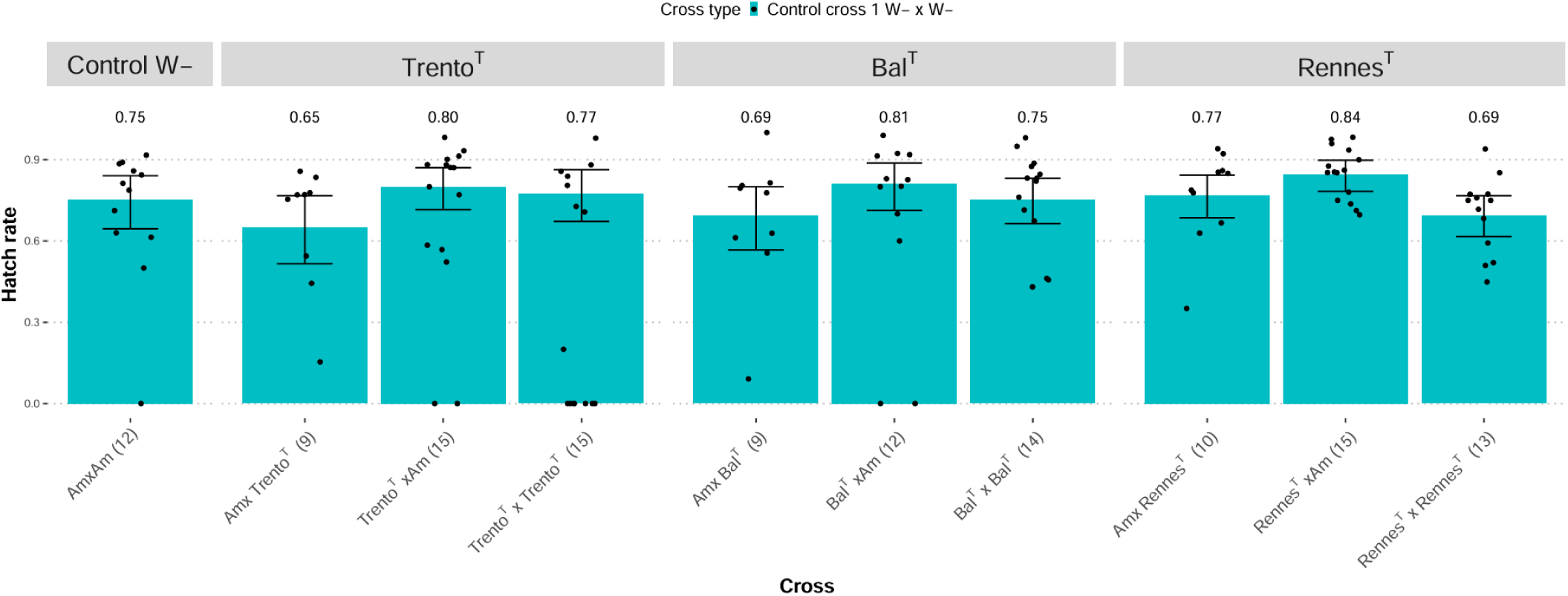
Hatch rates in crosses involving three tetracycline-treated *D. suzukii* lines (Trento^T^, Bal^T^, and Rennes^T^). Crosses are given as “female line × male line” (W+, *Wolbachia*-infected line; W−, uninfected line). After tetracycline treatment, lines were free of *Wolbachia*. Bars show fitted hatch rates excluding structural zeros, estimated from the zero-inflated beta-binomial model, with 95% confidence intervals obtained by parametric bootstrap (500 replicates); points show the observed hatch rate of each couple. The number of couples assayed is given in parentheses below each bar. The uninfected control cross (Am × Am) is shared by the three lines and is shown once, on the left.

### Appendix S3 New draft genome assembly of *w*Suz (*w*Suz_RW2025a) and phylogeny of *cif* genes

The previously available *w*Suz assembly (GCA_000333795.2; Siozios et al., 2013) is highly fragmented (110 contigs, totalling ∼1.35 Mb; largest contig 89,713 bp). This limits analyses of genome structure and of genomic regions of interest such as the *cif* loci. We therefore generated a new assembly, *w*Suz_RW2025a, with substantially improved contiguity.

We retrieved publicly available PacBio long-read data (SRA SRX27255168) generated by Wang et al. (2025) to assemble a reference *D. suzukii* genome from individuals collected in China (Wang et al., 2025). A preliminary *k*-mer-based screening (Gautier, 2023) indicated that these individuals were infected with *Wolbachia*. We then reassemble the whole-genome from these data using hifiasm with default parameters (v0.25.0; Cheng et al., 2021) and searched for *Wolbachia* contigs in the resulting primary assembly (consisting of 1,037 contigs totalling 282 Mb, N50=6.8Mb) by querying a database constructed from the NCBI nonredundant nucleotides (nt) released in February 2020 using the program Kraken2 (v2.1.2; Wood et al., 2019). This allowed identification of four candidate *w*Suz contigs (of 841,453; 270,644; 215,926 and 54,687 bp) totaling 1,382,710 bp.

To evaluate the gain in contiguity, we compared *w*Suz_RW2025a with the previous *w*Suz assembly. To assess its structure relative to a complete genome of a closely related strain, we also compared it with *w*Ri from

*D. simulans* (GCA_000022285.1; Klasson et al., 2009). Whole-genome alignments were computed and visualised with D-GENIES v1.5.0 using default parameters (Cabanettes & Klopp, 2018).

Numerous fragments of the previous *w*Suz assembly aligned along the larger contigs of *w*Suz_RW2025a, supporting the improved contiguity of the new assembly (Fig. S2B). Comparison with *w*Ri revealed extensive homologous regions distributed across all *w*Suz_RW2025a contigs, in both orientations relative to *w*Ri (Fig. S2A). This pattern is consistent with differences in contig order and orientation, structural rearrangements between the two genomes, or both.

We used BLAST searches with the Type I and Type II *cif* sequences of *w*Ri. To assess the phylogenetic position of the retrieved *w*Suz *cif* genes, we used the *cif* sequences of *w*Ri, *w*Mel and *w*Pip (Martinez et al., 2021). For each copy of a Type I or Type II *cif* pair, nucleotide sequences were translated with the Biostrings R package (Pagès et al., 2024). CifA and CifB were aligned separately with MAFFT (Katoh & Standley, 2013) and trimmed with ClipKIT (Steenwyk et al., 2020), both with default parameters, and the two alignments were then concatenated. A maximum-likelihood tree was inferred with IQ-TREE 3 (Wong et al., 2026), with the substitution model selected by ModelFinder (Kalyaanamoorthy et al., 2017) and branch support from 1,000 ultrafast bootstrap replicates. The tree was drawn with ggtree (Yu et al., 2017).

**Fig. S2.**
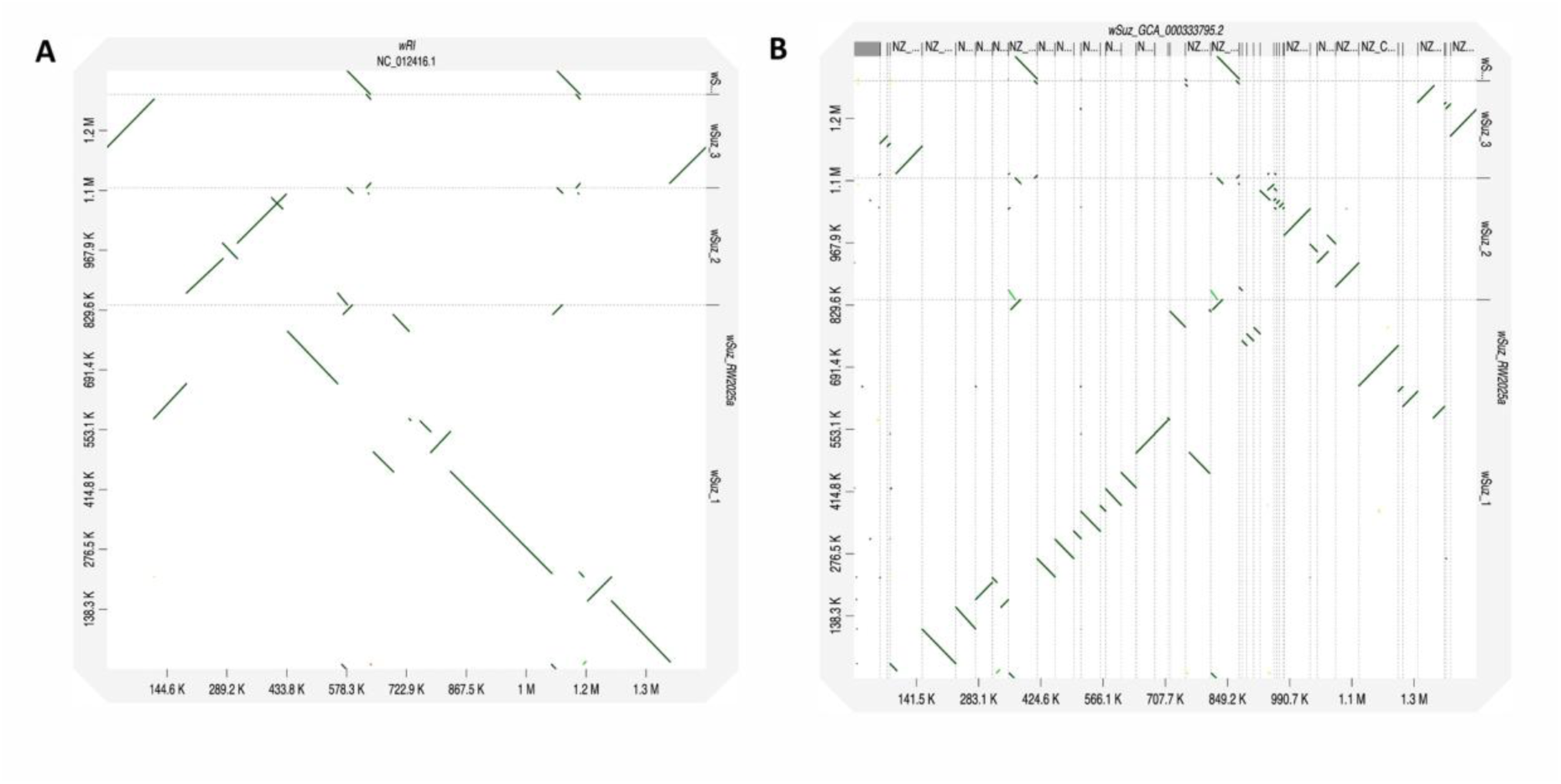
Dotplots generated with dgenies v1.5.0 (Cabanettes & Klopp, 2018) (default options) comparing the *w*Suz_RW2025a with (A) the *w*RI reference assembly (GenBank ID: GCA_000022285) of the closely related *Wolbachia* from *D. simulans* or (B) the previous and more fragmented *w*Suz assembly (GenBank ID: GCA_000333795.2).

### Appendix S4 Genetic diversity among *Wolbachia* isolates from the four lines of *D. suzukii*

To genotype *Wolbachia* isolates and verify the presence of and polymorphism in *cif* genes, we performed whole-genome sequencing of the four lines of *D. suzukii* (Bal, Par6, Rennes and Trento). DNA extraction was performed from ovaries to recover endosymbiont-enriched samples (Jousselin et al., 2016). Briefly, for each line, the ovaries of 25 females were isolated and preserved in 96% ethanol and sequentially filtered through two filters (with pores of 100 µm or 30 µm; SEFAR, Switzerland) and two Millipore Isopore^TM^ white polycarbonate membranes with 10-µm and 5-µm pores, respectively (Merck Millipore, Ireland). After filtration and centrifugation, we extracted DNA using the DNeasy Blood & Tissue Kit (Qiagen, USA) according to the manufacturer’s recommendations. DNA was mechanically fragmented to 300-600 bp fragments using a Bioruptor Pico (Diagenode; 15s ON, 90s OFF, eight cycles). Libraries were constructed using the NEBNext Ultra II DNA Library prep kit for Illumina (New England Biolabs) with IDT for Illumina TruSeq DNA UD Indexes (ref 20022370, Illumina, USA). All libraries were pooled at an equimolar ratio and paired-end sequenced (2 × 250 bp) on a NovaSeq 6000 sequencer (SP flow cell) at the GeT-PlaGe sequencing facility (INRAE, Toulouse, France). Raw sequencing data are available at the European Nucleotide Archive (ENA) under study accession PRJEB127904.

The SRA accessions used for phylogenomic analyses are listed in Table S10. We included *Wolbachia* WGS data used to assemble the reference genome of *w*Suz from Trento, Italy (Siozios et al., 2013) and from *D. subpulchrella* or *D. simulans* individuals infected with either *w*Spc or *w*Ri. WGS reads were first filtered using *fastp* (v0.23.2; Chen et al., 2018) run with default options. Filtered reads were mapped onto our *w*Suz reference genome (Zenodo, https://doi.org/10.5281/zenodo.24090511), complemented with the complete mitochondrial sequence of *D. suzukii* (GenBank Accession ID: KU588141.1) and the reference genome of *D. suzukii* (GenBank Accession ID: GCA_043229965.1; Camus et al., 2025). Read mapping, removal of PCR duplicates and joint variant calling across all samples followed the pipeline of Camus et al., 2025, with bwa-mem2 v2.2.1, SAMtools v1.19 and FreeBayes v1.3.10. We filtered out variant positions found in less than 90% of the ten *Wolbachia* genomes using the R package poolfstat (v3.1; Gautier et al., 2022). To compute the genetic distance matrix, we computed pairwise divergence between pairs of isolates using the compute.fstats function. The resulting pairwise divergence matrix was used to infer a neighbor-joining (NJ) phylogenetic tree with the nj function from the ape package (Paradis & Schliep, 2019). The robustness of the inferred phylogenetic relationships was assessed by bootstrap resampling of the variant dataset. For each of 1,000 bootstrap replicates, variants were sampled with replacement, pairwise nucleotide divergence was recalculated using poolfstat, and a corresponding NJ tree was reconstructed. Bootstrap support for each branch was calculated as the percentage of bootstrap trees recovering the corresponding split. The final NJ tree and bootstrap support values were visualized and annotated using ggtree in R.

To estimate the copy number of *cif* genes in the *w*Suz genome, we compared their read depth with that of single-copy genes. Per-base read depth of primary alignments across the *w*Suz reference genome was calculated from the deduplicated BAM files using SAMtools (-F 0×900 flag; Li et al., 2009). For each gene, we computed per-base read depth normalized by the mean depth of its contig, and reported the mean and SD of this normalized depth across gene positions.

To verify that normalized read depth recovers the known copy number of *cif* genes, we applied the same approach to two sequenced isolates of *w*Ri (*w*Ri US-CA and wRi ZB; Turelli et al., 2018), a *Wolbachia* strain in which the number of *cif* loci has been established from complete genome assemblies (Klasson et al., 2009). *w*Ri carries two Type I and one Type II *cifA–cifB* pairs. Normalized read depth recovered these copy numbers in both isolates: Type I *cifA* and *cifB* reached approximately twice the depth of the single-copy genes *ftsZ* and *wsp*, whereas Type II genes were at single-copy depth (Fig. S3). This confirms that normalized read depth provides a reliable estimate of *cif* copy number in *w*Suz isolates (Fig. 4D).

**Fig S3.**
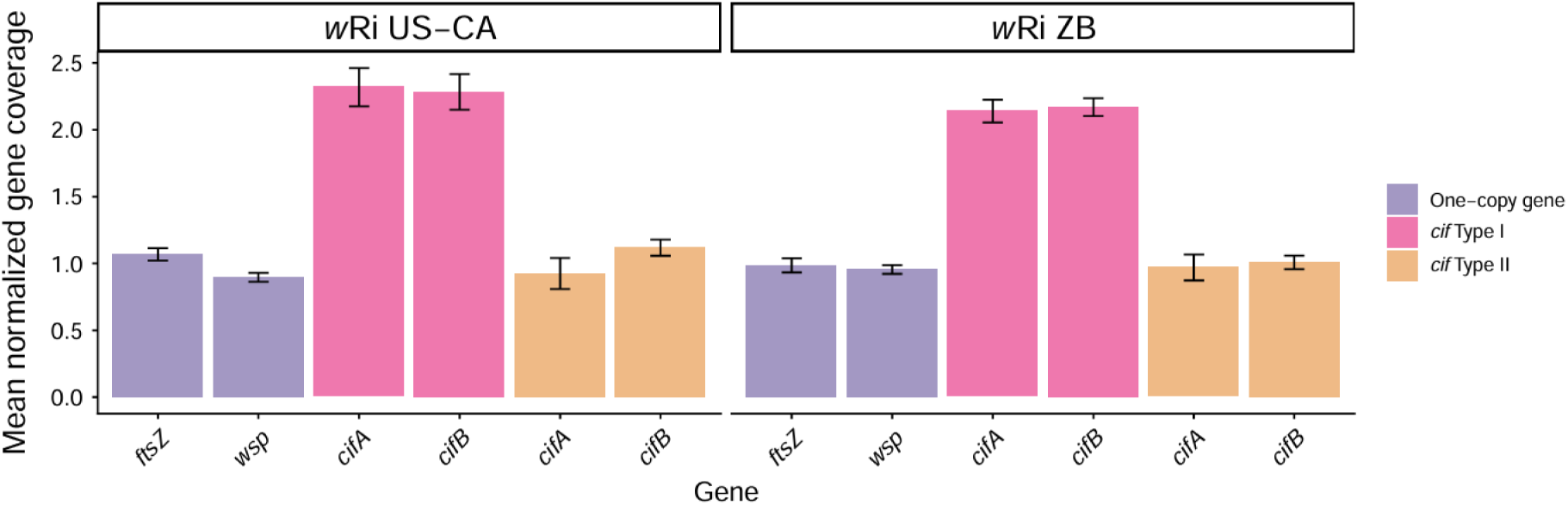
Normalized gene coverage in two sequenced isolates of *w*Ri. Mean normalized read depth of the single-copy genes *ftsZ* and *wsp*, and of the Type I and Type II *cifA* and *cifB* genes, in *w*Ri US-CA and *w*Ri ZB. Per-base read depth was normalized by the mean depth of the corresponding contig, as described above; bars show the mean and error bars show the SD of normalized depth across gene positions.

### Appendix S5 Females infected with *Wolbachia* rescue CI in *D. suzukii*

To test whether *Wolbachia*-infected females rescue the reduction in hatch rate caused by infected males, we used males from the two lines in which CI was detected, Bal and Trento. Males of each line were crossed with uninfected Am females (CI cross, W⁻ × W⁺) and with infected females of their own line and of other infected lines (rescue crosses, W⁺ × W⁺): Bal, Trento and Par6 females with Bal males, and Trento and Bal females with Trento males. Par6 and Rennes males were not included because CI was undetected in these lines (Table S2c).

Hatch rates were measured as described in Materials and Methods, in two experimental blocks (batch 2 and 4). For each male line, hatch rates were analysed with zero-inflated beta-binomial models, with a constant zero-inflation term and a common dispersion parameter. We compared a null model (no difference among test and control crosses), a model contrasting the hatch rate in the test cross vs. rescue crosses with infected females, and a saturated model with a separate hatch rate for each cross. CI strength was estimated as the relative reduction in hatch rate of the CI cross compared with each rescue cross. 95% confidence intervals were obtained by parametric bootstrap with 500 iterations.

For both male lines, the null model received little support (Bal: ΔAICc = 20.43; Trento: ΔAICc = 10.13; Table S5a; Fig.S4). With Bal males, infected females from different lines differed in hatch rate: the saturated model had lower AICc than the model contrasting only the CI cross with the rescue crosses (ΔAICc = 3.55; Table S5a). These differences may reflect differences in baseline hatch rate among female lines or could reflect differences in the ability to rescue CI. With Trento males, the two models could not be distinguished (ΔAICc = 0.80).

These results indicate efficient rescue of CI by *Wolbachia*-infected females: in all line combinations tested, hatch rate in rescue crosses was higher than in the CI cross with the same males (Table S5b,c)

**Fig. S4.**
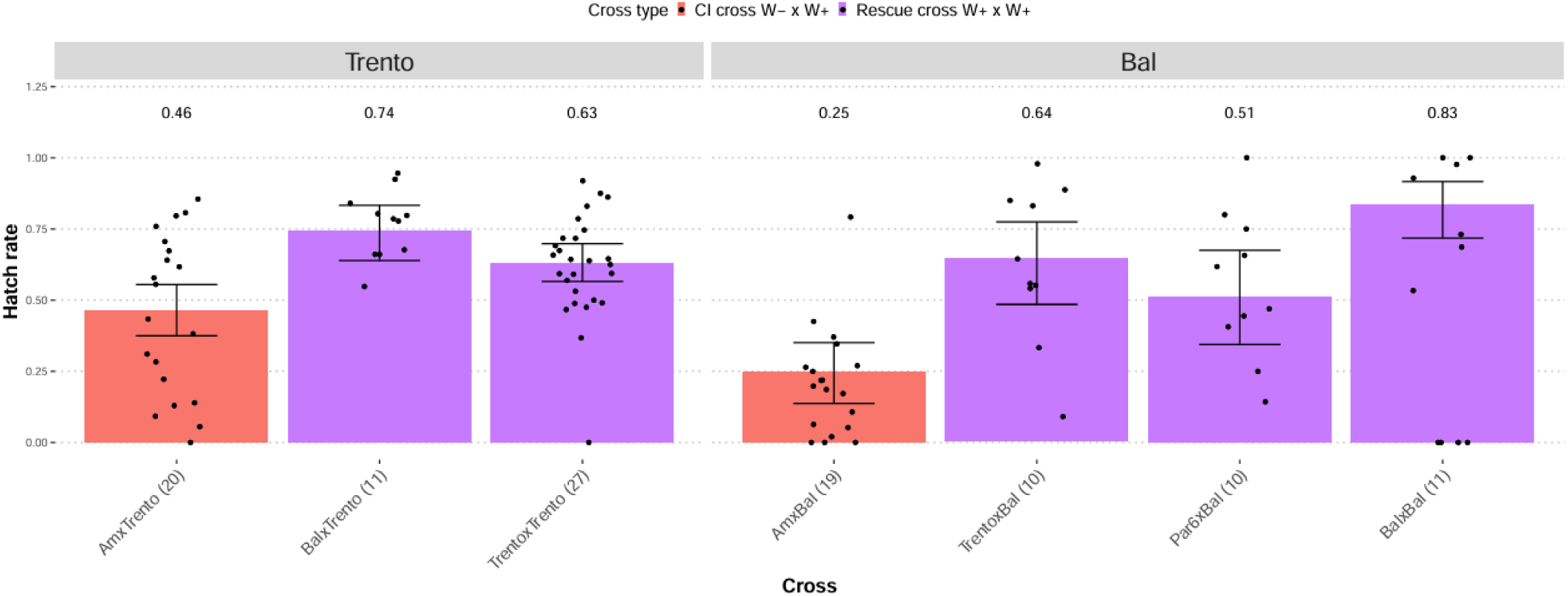
Hatch rates in CI and rescue crosses between *D. suzukii* lines naturally infected with *Wolbachia*. Crosses between uninfected Am females and infected Bal or Trento males were used as controls for CI occurrence (cross W− × W+); rescue was tested in crosses between infected Bal or Trento males and infected females of their own or of other infected lines (cross W+ × W+). Crosses are given as “female line × male line” (W+, infected line; W−, uninfected line). Bars show fitted hatch rates excluding structural zeros, estimated from the beta-binomial model, with 95% confidence intervals obtained by parametric bootstrap (500 replicates); points show the observed hatch rate of each couple. The number of mating pairs assayed is given in parentheses below each bar.

### Appendix S6 *w*Suz isolates do not induce detectable CI in the Am male host nuclear background

To test whether the host nuclear background affects CI expression, we introgressed the cytoplasm, and thus the *Wolbachia*, of the lines inducing strong CI (Bal and Trento) and of the lines with weak or undetected CI (Rennes and Par6) into a single host nuclear background, that of the initially uninfected Am line. To do introgression, we made seven generations of backcrosses of females from a *Wolbachia*-infected line with males of the Am strain. We obtained Am^Par6^, Am^Rennes^, Am^Trento^ and Am^Bal^ lines of *D. suzukii* with four different cytoplasms but an almost identical nuclear genetic background (more than 99% of the Am genetic background).

To test whether the host nuclear background affects the amount of *Wolbachia* within the host, we estimated the *Wolbachia* genome copy number relative to the host genome copy number in each introgressed line using TaqMan qPCR (Cattel, Martinez, et al., 2016). To standardize male developmental conditions, ten females and two males were placed in each of three tubes. Females were allowed to lay eggs for two days before adults were removed. Emerging male flies were collected daily and transferred to new tubes, where they were aged for three days. Male testes were dissected on ice in a PBS solution (BioBasic, Canada). For each line, five pairs of testes from males originating from the same tube were pooled together in a 1.5 mL tube containing 200 µl of ATL lysis buffer (Qiagen, USA) and stored at −80°C until DNA extraction. We replicated the process five times (five pools × five testes pairs per line).

DNA was extracted using a DNeasy Blood & Tissue kit (Qiagen, USA). qPCR was performed following the protocol described in Cattel, Martinez, et al., 2016, with modifications. The *Wolbachia* genome copy number was quantified using *ftsZ* primers (FtsZ_F157: 5’-CGA-GAT-GGG-CAA-AGC-GAT-GA-3’; FtsZ_R282: 5’-ATT-CCT-TGC-GCA-CCT-TTC-AT-3’) with the probe FtsZ_P182 (6-FAM-ACC-GGA-GAG-GCA-GAA-GGA-GA-BHQ-1), while the host genome was quantified using the single-copy *Rpl32* gene (primers: rp49Fdd: 5’-CTG-CCC-ACC-GGA-TTC-AAG-3’; rp49Rdd: 5’-CGA-TCT-CGC-CGC-AGT-AAA-C-3’; probe RP49dd_P HEX-CCT-CCA-GCT-CGC-GCA-CGT-TG-BHQ-1). PCR reactions were conducted on a LightCycler 480 Real-Time PCR System (Roche, Basel, Switzerland) with two technical replicates per sample. Samples were distributed across a 384-well plate. We generated efficiency curves using a series of dilutions of purified PCR products obtained from infected *D.suzukii* DNA samples. Two types of negative controls were included: DNA from two *Wolbachia-*uninfected lines and a template-free control containing deionized water. Each reaction was performed in 10 µl reaction volume consisting of 5 µl 2X LightCycler 480 Probes Master Mix (Roche, Basel, Switzerland), 0.45 µM of each primer and 0.45 µM of probe and 2 µl DNA sample (0.3-1.5 ng/µl). The following cycling conditions were used for amplification: initial preincubation 10 min at 95°C, followed by 50 cycles of 15 s at 95°C, 30 s at 60°C, and 1 s at 72°C, with final cooling 10 min at 40°C. Fluorescence detection was performed during the step at 60°C of each amplification cycle. All Ct values used for *Wolbachia* titer estimation were below 35.

Wolbachia titer was calculated based on C_t_ value as described in Hague et al., 2020 using the 2^ΔCt^ method (Livak & Schmittgen, 2001). Briefly, we calculated the mean cycle threshold (C_t_) value for two replicates for each sample and estimated *Wolbachia* titer with the formula:

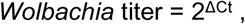

where ΔC_t_ = C_t*Host*_− C_t*Wolb*_ *, C_tWolb_* and *C_tHost_* represent cycle threshold values for the *Wolbachia ftsZ* and Drosophila *Rpl32* amplification, respectively.

*Wolbachia* titers were log_10_–transformed to meet the assumptions of normality and homoscedasticity, verified by inspection of residual and quantile–quantile plots, and compared among lines by ANOVA followed by Tukey-adjusted pairwise contrasts on the log scale (R package emmeans; Lenth, 2024). Ratios of titers between lines and their 95% confidence intervals were obtained by back-transformation.

Titers did not differ detectably among the four introgressed lines (ANOVA on log₁₀-transformed titers: F_3,16_ = 1.55, P = 0.24), regardless of their cytoplasmic origin (Trento, Bal, Par6, or Rennes), with line means ranging from 17.8 to 25.6 *Wolbachia* genome copies per host genome copy (Fig. S5). In contrast, the corresponding naturally infected lines differed substantially in titer (F_3,16_ = 6.17, P = 0.0055; means from 8.0 to 48.2 copies per host genome copy; Fig. 3). These results suggest that the host nuclear background contributes to variation in *Wolbachia* titer.

**Fig. S5.**
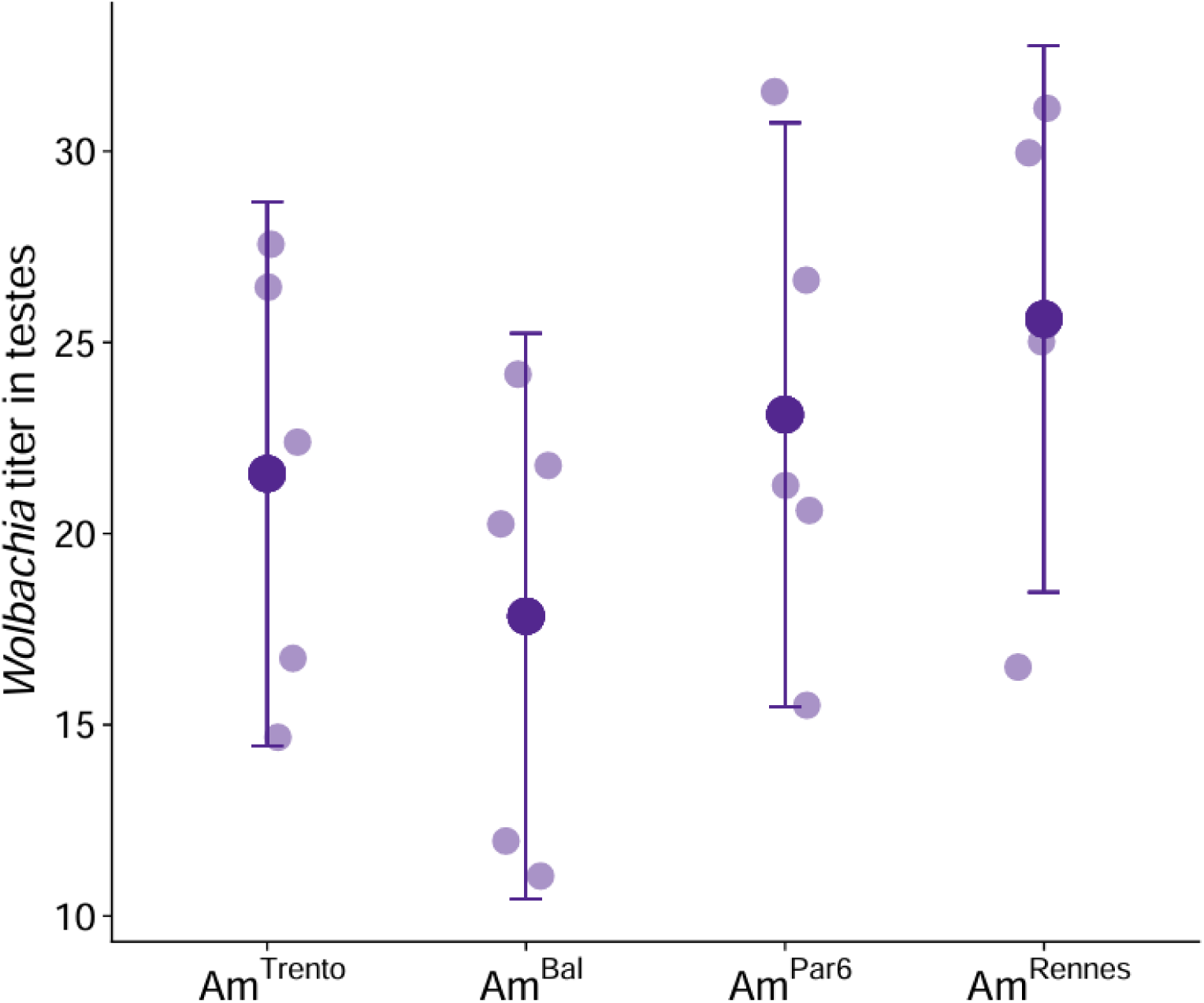
*Wolbachia* titers in testes of 3-day-old *D. suzukii* males carrying the *Wolbachia* isolates of Trento, Bal, Par6 and Rennes in the Am nuclear background (Am^Trento^, Am^Bal^, Am^Par6^, Am^Rennes^). Titer was calculated as 2^ΔCt^ of the *Drosophila* gene *RpL32* to the *Wolbachia* gene *ftsZ* and is expressed as the number of *Wolbachia* genome copies per host genome copy (n = 5 samples per line). Small circles show individual samples; large circles and bars show means and 95% confidence intervals. None of these lines induced detectable CI. Titers did not differ among lines (ANOVA on log₁₀-transformed titers: F_3,16_= 1.55, P = 0.24).

We conducted the CI assays of introgressed lines (Am^Par6^, Am^Rennes^, Am^Trento^, Am^Bal^) in two experimental blocks (blocks 6 and 7) following the procedure described in the main text. In the introgressed lines, CI was no longer detected. Hatch rates of the test cross (W− × W+) were similar to those of the three control crosses of the same line with overlapping confidence intervals (Fig. S6, Table S6b). Estimated CI strength relative to the uninfected control cross (95% confidence interval) was 9.2% (0–22.7%) in Am^Trento^, 9.2% (0–20.5%) in Am^Bal^, 4.5% (0–17.7%) in Am^Rennes^ and 1.2% (0–11.3%) in Am^Pa6^ (Table S6c). Model selection pointed the same way: the null model, with no difference among cross types, ranked first in Am^Bal^, Am^Rennes^ and Am^Par6^, and in Am^Trento^ the model contrasting the test cross with the control crosses was indistinguishable from the null model (ΔAICc = 0.23; Table S6a). By comparison, the naturally infected Trento and Bal lines showed strong CI, with CI strengths of 46.7% (38.3–54.4%) and 66.5% (57.6%–74.8%) (Table S2c).

Because the introgressed lines carry the same *Wolbachia* and cytoplasm as the naturally infected lines but the nuclear background of the uninfected Am line, the loss of CI in Am^Trento^ and Am^Bal^ indicates that the Am nuclear background is sufficient to suppress CI expression in these lines. Moreover, the four introgressed lines, which carry four *Wolbachia* isolates, did not induce detectable CI, whereas the naturally infected lines ranged from strong CI (Bal, Trento) to undetectable CI (Rennes, Par6). The variation in CI strength among lines is therefore determined by the nuclear genotype of the male rather than by the *Wolbachia* genotype.

**Fig. S6.**
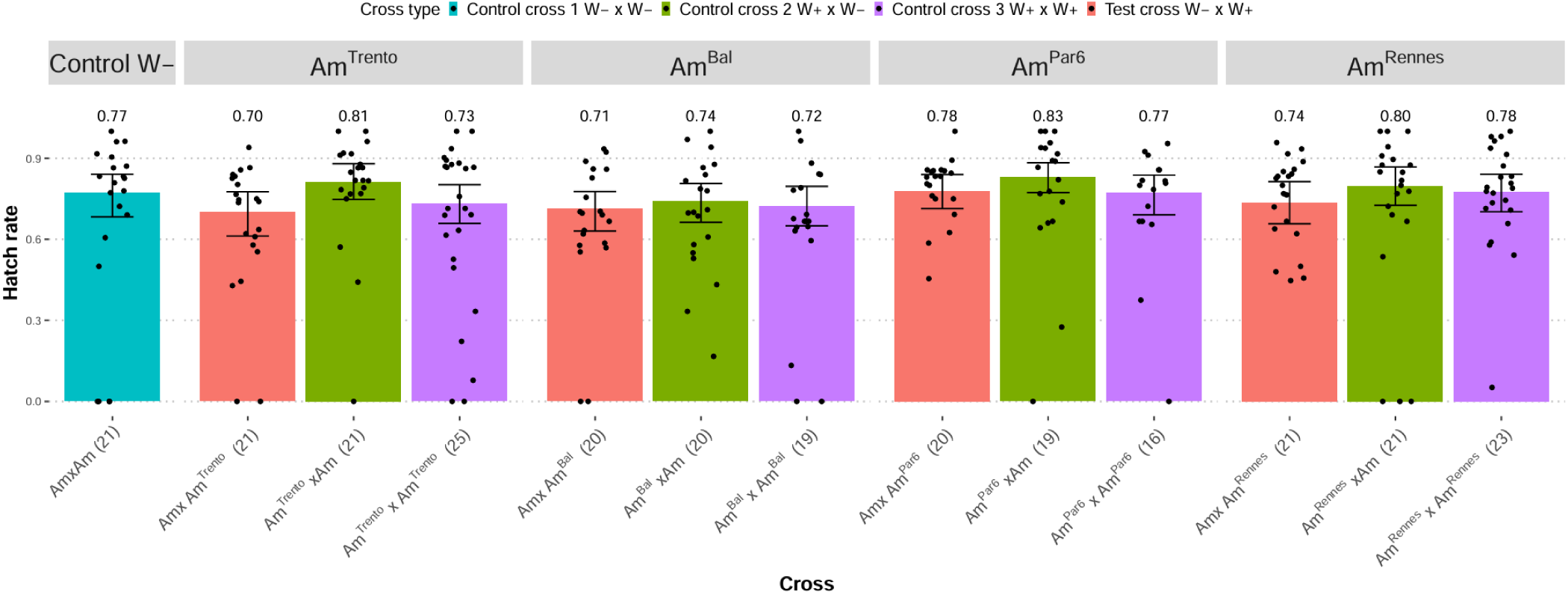
Hatch rate in CI assay for *D. suzukii* lines introgressed with four different *Wolbachia* isolates (Am^Trento^, Am^Bal^, Am^Par6^, Am^Rennes^). Crosses are given as “female line × male line” (W+, infected line; W−, uninfected line). Bars show fitted hatch rates excluding structural zeros, estimated from a beta-binomial model, with 95% confidence intervals obtained by parametric bootstrap (500 replicates); points show the observed hatch rate of each couple. The number of mating pairs assayed is given in parentheses below each bar.

### Appendix S7 Variation in *Wolbachia*-induced cytoplasmic incompatibility among male and female genetic backgrounds

#### Male genetic effect on CI

To test whether differences in hatch rate were associated with additive autosomal, dominance autosomal, X-linked or Y-linked genetic effects, we decomposed the differences in hatching success between the six crosses (Am females mated with Bal, Par6, F1 BalxPar6, F1 Par6xBal, F2 (BalxPar6)x(BalxPar6) or F2 (Par6xBal)x(Par6xBal) males), using the generation mean approach (Gilchrist & Partridge, 1999). Each male genotype was described by four genetic coefficients (Table 1 in Gilchrist & Partridge, 1999):

- Additive autosomal effect (*a*): the expected proportion of Par6 autosomal alleles, scaled from −1 (Bal) to +1 (Par6). It equals 0 in F1 and F2 males.
- Autosomal dominance effect (*d*): the expected proportion of heterozygous autosomal loci. It equals 0 in parental males, 1 in F1 males and 0.5 in F2 males.
- X-chromosome effect (*x*): +1 when the male’s X chromosome originates from Par6 and −1 when it originates from Bal. F2 males received their X from a heterozygous F1 mother, so they were assigned the expected value 0.
- Y-chromosome effect (*y*): +1 when the male’s Y chromosome originates from Par6 and −1 when it originates from Bal.

Hatch rate was modelled with a zero-inflated beta-binomial Generalised Linear Mixed Model using the package glmmTMB v1.1.15.2 (Brooks et al., 2017). For couple *i*, the number of hatched larvae *Y_ij_* out of *n_ij_* eggs was modelled using a Zero-inflated beta binomial distribution as:

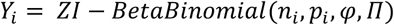

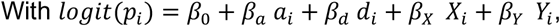

were π represents the probability of a structural zero (a couple with no offsrping larvae beyond that expected under the beta-binomial), *p_ij_* represents the expected hatch rate excluding structural zeros, φ the beta-binomial dispersion parameter 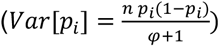.

Starting from the full model (logit(p)=*a* + *d* + *x + y*), we fitted a set of 15 nested models: the full model, all four three-term models, five two-term models (logit(p)=*a* + *d*, *a* + *x*, *a* + *y*, *d* + *x*, *d* + *y* or *x* + *y*) and the intercept-only model and the saturated model (with one parameter for the 6 types of crosses). All models shared the same dispersion and zero-inflation structure. They were ranked by their corrected Akaike’s information criterion (Hurvich & Tsai, 1989) using MuMIn v1.48.19 (Bartoń, 2026). Models within 2 AICc units of the best model were considered equally supported. Convergence was assessed by checking that the Hessian matrix was positive-definite and that log-likelihoods were consistent across nested models.

#### Parameter coefficients of the model for comparing hatch rate in test cross with males from Bal and Par6 (parental), F1 or F2 lines

| **Male genotype** | **Genetic effects** |  |  |  |
|---|---|---|---|---|
|  | **Additive autosomal** | **Dominance autosomal** | **X chromosome\*** | **Y chromosome\*** |
| Parental line 1 (Par6) | 1 | 0 | 1 | 1 |
| Parental line 2 (Bal) | -1 | 0 | -1 | -1 |
| F<sub>1</sub> Par6 x Bal | 0 | 1 | 1 | -1 |
| F<sub>1</sub> Bal x Par6 | 0 | 1 | -1 | 1 |
| F<sub>2</sub> (Par6 x Bal) x (Par6 x Bal) | 0 | 0.5 | 0 | -1 |
| F<sub>2</sub> (Bal x Par6) x (Bal x Par6) | 0 | 0.5 | 0 | 1 |

\*Only F<sub>2</sub> males allow disentangling Y-linked from X-linked genetic effects, as among males with a Par6 grandfather, 100% have a Par6 Y chromosome, but 50% have a X chromosome from Par6 and the other 50% have a X chromosome from Bal.

#### Number of suppressor loci

The minimum effective number of loci underlying CI suppression was estimated with the Castle–Wright estimator, 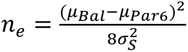 (Castle, 1921; Lande, 1981). Here *μ_Bal_* and *μ_Par_*_6_ are the logit of the mean hatch rates of the parental male genotypes, and *σ*^2^ is the segregation variance, estimated as the between-male variance among F2 males minus that among parental and F1 males. Analyses were based on egg and larval counts from each Petri dish (two to three dishes per mating pair), rather than on counts pooled per pair. Estimates were obtained on the logit scale from a zero-inflated beta-binomial mixed model with cross type as a fixed effect and separate between-male (couple) variances for F2 and non-F2 males. Confidence intervals were obtained by parametric bootstrap (500 simulations). Refits that failed or yielded a non-positive-definite Hessian (7.6%) were discarded. Because *n*ₑ is undefined when the estimated segregation variance is zero or negative, we computed percentile intervals for 1/*n*ₑ, which is always finite, and inverted them to obtain intervals for *n*ₑ. Between-male variance in hatch rate (logit scale) did not differ among Bal, Par6 and F1 males (likelihood-ratio test, χ² = 0.02, df = 2, *P* = 0.99). This supports the assumption of homogeneous environmental variance underlying the Castle–Wright estimator, and these groups were pooled to estimate it. Between-male variation in hatch rate for each cross type is shown in Fig. S7.

**Fig. S7.**
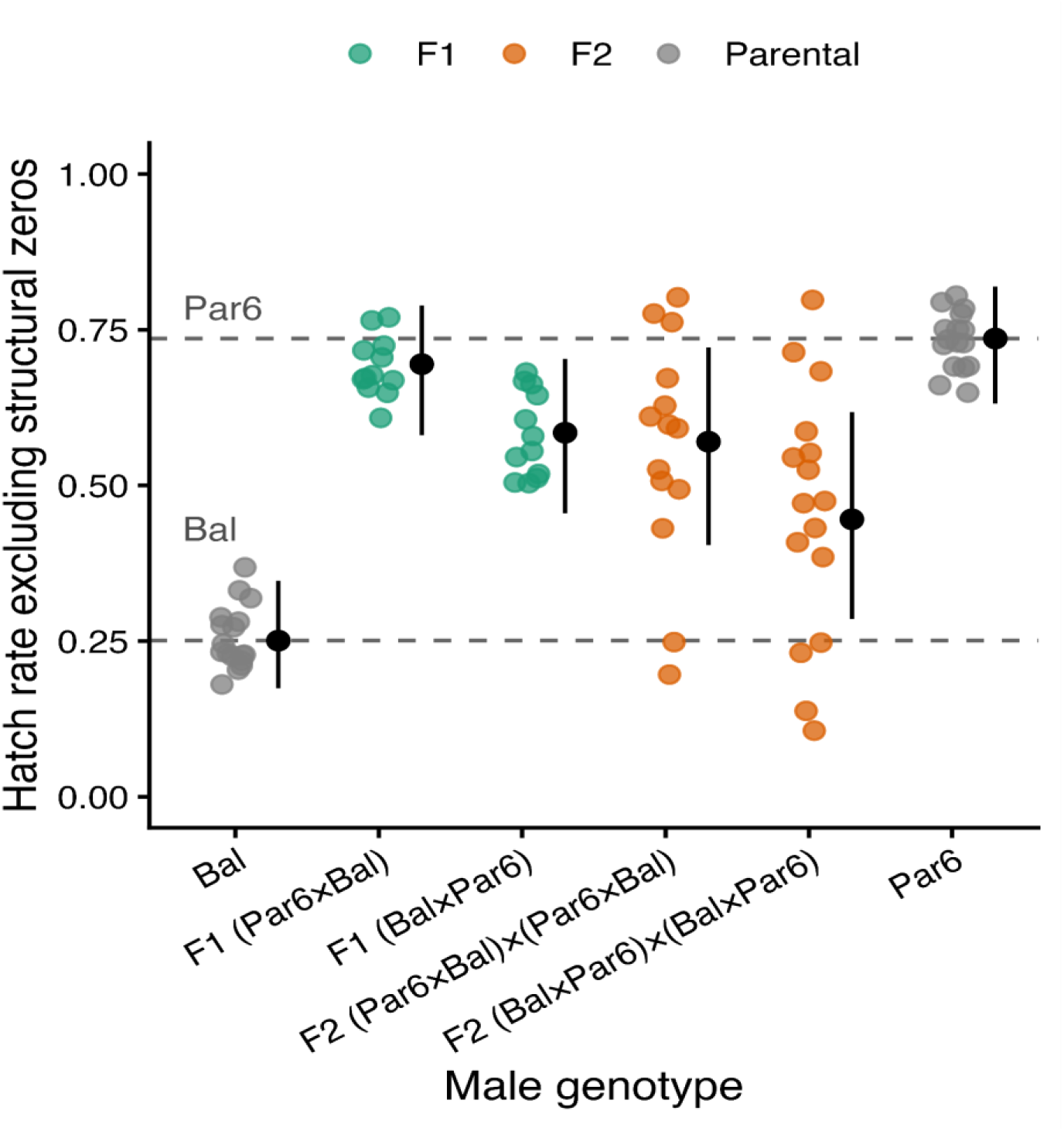
Hatch rates of embryos from Am females mated with parental, F1 or F2 males. Each colored point represents the predicted hatch rate of an embryo derived for each male in the dataset (n = 87 males, 2–3 Petri dishes each) based on a zero-inflated beta-binomial generalized linear mixed model with cross type as a fixed effect and separate between-male variances for F2 and non-F2 males. Hatch rates exclude structural zeros (dishes with no hatching attributed to the zero-inflation component). Black points and bars show the mean hatch rate and 95% confidence interval for each cross type (typical couple random effect set to zero). Dashed lines indicate the mean hatch rates with Bal and Par6 males.

#### No variation in CI among uninfected females of different genotypes

To test whether CI induced by infected males differs among the genetic backgrounds of uninfected females, we crossed males from Bal with females from two additional uninfected strains (Mon47 and Par7), while we crossed Par6 males with females from the uninfected line Par7 and Rennes males with females from the Mon47 uninfected line. To assay CI, we used the same protocol as the one described in Materials and Methods. For each of the three male lines, we tested for differences in hatch rates among crosses with different female genotypes. For each male line, the response was the proportion of eggs that hatched per mating pair, weighted by the number of eggs laid. The conditional (beta-binomial) part of the model tests for differences in hatch rate among female genotypes. We fitted two models (Table S8). (i) In the null model, the mean hatch rate was constant across female genotypes. (ii) In the female genotypes model, the mean hatch rate differed among female genotypes. All models included a constant zero-inflation parameter (accounting for variation among pairs in the probability of producing no larvae) and a constant dispersion parameter (accounting for overdispersion in hatch rate among mating pairs).

For Bal, Rennes or Par6 males, hatch rates in test crosses were similar across the female backgrounds tested and the female genotype model had the same log-likelihood as the null model (Fig. S8; Tables S8a and S8b). Thus, female genotype had no detectable effect on hatch rate.

**Fig. S8.**
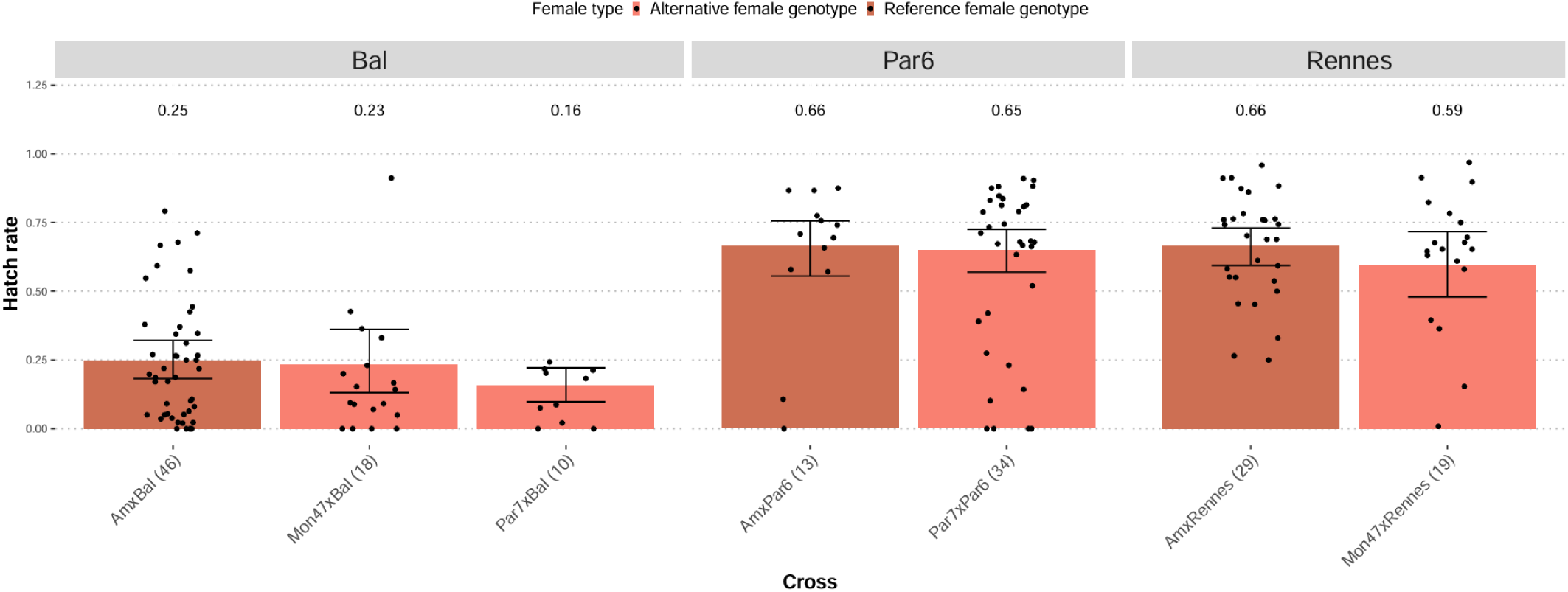
Hatch rates in test crosses with different uninfected female genotypes. Test crosses were performed between males from the *Wolbachia*-infected Bal, Par6, and Rennes lines and females from different uninfected *D. suzukii* lines. Crosses are given as “female line × male line” (W+, infected line; W−, uninfected line). Bars show fitted hatch rates, estimated from the saturated model, with 95% confidence intervals obtained by parametric bootstrap (500 replicates); points show the observed hatch rate of each couple. The number of mating pairs assayed is given in parentheses below each bar.

### Appendix S8: Male age between 2 and 3 days does not affect CI expression

Males used in our CI assays were 2–3 days old at the time of mating. To test whether variation within this age range influenced CI expression, we fitted zero-inflated beta-binomial models with the same specification as for the CI assay (constant zero-inflation and dispersion; Materials and Methods) with male line as a fixed effect on hatch rate, and compared them with models that additionally included male age as a continuous explanatory variable. We analysed three data sets: test crosses between uninfected females and infected males (W⁻ × W⁺) of the four lines (n = 129 pairs); the same test crosses restricted to the two lines with strong CI, Bal and Trento (n = 87); and Control cross 3 between infected females and males of the same line (W⁺ × W⁺; n = 127), in which CI is rescued, and any age effect reflects male fertility rather than CI. Par6 males in test crosses were all 3 days old and therefore did not contribute to the estimate of the age effect in that data set. Models with and without male age were compared using AICc. The age effect was estimated from the model including male age and is reported as the odds ratio of hatching per additional day of male age, with 95% confidence intervals obtained by parametric bootstrap (500 replicates; Table S9a-c).

In all three analyses, the model without male age had the lower AICc (test crosses, all lines: ΔAICc = 2.18; test crosses, Bal and Trento: ΔAICc = 2.21; Control cross 3: ΔAICc = 1.90; Table S9a). The confidence intervals for the age effect included 1 in all three analyses (odds ratio per day: test crosses, all lines, 1.06, 95% CI 0.69– 1.71; test crosses, Bal and Trento, 0.94, 0.54–1.81; Control cross 3, 1.14, 0.73–1.72; Table S9c), and estimated hatch rates changed little between 2- and 3-day-old males in any line (Table S9b). Thus, no effect of male age on hatch rate was detected within the 2–3 day range, either in test crosses or in compatible crosses. Given the width of the confidence intervals, moderate age effects cannot be excluded, but variation in male age within this narrow range is unlikely to account for the differences in CI strength among lines.

**Table S1.** List of the live lines of *D. suzukii* used for this study. Wolbachia infection status was determined by PCR amplification of ftsZ and wsp, Drosophila D14 locus was used as a control; wSuz-infection-consensus gives the status inferred from the two markers.

| Line_ID | Town | Country | GPS coordinates (latitude, longitude) | Sampling date | Collector | Line type | Number of individuals tested | Type of material | *D14* | *ftsZ* | *wsp* | wSuz-infection-consensus | Used in this study | Reference |
|---|---|---|---|---|---|---|---|---|---|---|---|---|---|---|
| Am | Sospel | France | 43.88, 7.44 | July 2011 | Nicolas Gompel | Isofemale line | 10 | live flies | positive | negative | negative | uninfected | yes | Karageorgi et al 2017 |
| Par2 | Paris | France | 48.84, 2.36 | October 2014 | Antoine Fraimout | Isofemale line | 5 | flies in ethanol | positive | negative | negative | uninfected | no | Fraimout et al 2017 |
| Par6 | Paris | France | 48.84, 2.36 | October 2014 | Antoine Fraimout | Isofemale line | 5 | flies in ethanol | positive | positive | positive | infected | yes | Fraimout et al 2017 |
| Par7 | Paris | France | 48.84, 2.36 | October 2014 | Antoine Fraimout | Isofemale line | 5 | flies in ethanol | positive | negative | negative | uninfected | yes | Fraimout et al 2017 |
| Par8 | Paris | France | 48.84, 2.36 | October 2014 | Antoine Fraimout | Isofemale line | 10 | flies in ethanol | positive | negative | poorly visible band | uninfected | no | Fraimout et al 2017 |
| Par9 | Paris | France | 48.84, 2.36 | October 2014 | Antoine Fraimout | Isofemale line | 10 | flies in ethanol | positive | negative | poorly visible band | uninfected | no | Fraimout et al 2017 |
| Par21 | Paris | France | 48.84, 2.36 | October 2014 | Antoine Fraimout | Isofemale line | 5 | flies in ethanol | positive | negative | negative | uninfected | no | Fraimout et al 2017 |
| Par22 | Paris | France | 48.84, 2.36 | October 2014 | Antoine Fraimout | Isofemale line | 5 | flies in ethanol | positive | negative | negative | uninfected | no | Fraimout et al 2017 |
| Par26 | Paris | France | 48.84, 2.36 | October 2014 | Antoine Fraimout | Isofemale line | 5 | flies in ethanol | positive | negative | negative | uninfected | no | Fraimout et al 2017 |
| Mon15 | Prades-le-Lez | France | 43.70, 3.87 | September 2015 | Anne Xuéreb | Isofemale line | 5 | flies in ethanol | positive | negative | negative | uninfected | no | Fraimout et al 2017 |
| Mon20 | Prades-le-Lez | France | 43.70, 3.87 | September 2015 | Anne Xuéreb | Isofemale line | 5 | flies in ethanol | positive | negative | negative | uninfected | no | Fraimout et al 2017 |
| Mon25 | Prades-le-Lez | France | 43.70, 3.87 | September 2015 | Anne Xuéreb | Isofemale line | 5 | flies in ethanol | positive | negative | negative | uninfected | no | Fraimout et al 2017 |
| Mon47 | Prades-le-Lez | France | 43.70, 3.87 | September 2015 | Anne Xuéreb | Isofemale line | 10 | flies in ethanol | positive | negative | negative | uninfected | yes | Fraimout et al 2017 |
| Bal | Bellegarde | France | 43.75, 4.46 | 2013 | Unknown | Lab colony | 5 | live flies | positive | positive | positive | infected | yes | Taning et al 2016 |
| West | Westland | Netherlands | Unknown | 2018 | Bregje Wertheim | Lab colony | 5 | live flies | positive | negative | negative | uninfected | no | Kruitwagen et al 2021 |
| Winssen | Beuningen | Netherlands | 51.89, 5.69 | 2018 | Bregje Wertheim | Lab colony | 5 | live flies | positive | negative | negative | uninfected | no | Gurung et al 2023 |
| Trento IAEA | Trento province | Italy | Unknown | Before 2013 | Omar Rota-Stabelli | Lab colony | 5 | live flies | positive | positive | positive | infected | no | Nikolouli et al 2020 |
| Rennes | Thorigné-Fouillard | France | 48.15, 1.58 | September 2017 | Hervé Colinet | Lab colony made out of nine isofemale lines | 10 | live flies | positive | positive | positive | infected | yes | Grumiaux et al 2019 |

**References**
Fraimout A, Debat V, Fellous S, et al (2017) Deciphering the routes of invasion of Drosophila suzukii by means of ABC random forest. Mol Biol Evol, 34:msx050.
Gurung, K., Vink, S. N., Salles, J. F., & Wertheim, B. (2023). More persistent bacterial than fungal associations in the microbiota of a pest insect. Journal of Pest Science, 96(2), 785-796.
Grumiaux, C., Andersen, M. K., Colinet, H., & Overgaard, J. (2019). Fluctuating thermal regime preserves physiological homeostasis and reproductive capacity in *Drosophila suzukii*. Journal of Insect Physiology, 113, 33-41.
Kruitwagen, A., Wertheim, B., & Beukeboom, L. W. (2021). Artificial selection for nonreproductive host killing in a native parasitoid on the invasive pest, *Drosophila suzukii*. Evolutionary Applications, 14(8), 1993-2011.
Karageorgi, M., Bräcker, L. B., Lebreton, S., Minervino, C., Cavey, M., Siju, K. P., ... & Prud'homme, B. (2017). Evolution of multiple sensory systems drives novel egg-laying behavior in the fruit pest *Drosophila suzukii*. Current Biology, 27(6), 847-853.
Nikolouli, K., Sassù, F., Mouton, L., Stauffer, C., & Bourtzis, K. (2020). Combining sterile and incompatible insect techniques for the population suppression of *Drosophila suzukii*. Journal of pest science, 93(2), 647-661.
Taning, C. N. T., Christiaens, O., Berkvens, N., Casteels, H., Maes, M., & Smagghe, G. (2016). Oral RNAi to control *Drosophila suzukii*: laboratory testing against larval and adult stages. Journal of pest science, 89(3), 803-814.

**Table S2a.** Results of the AICc model selection for hatch rate in four European *D. suzukii* lines in CI assay. Models are ranked from lowest to highest AICc values, and the first model listed for each strain corresponds to the best-supported model.

| Line | Model | Hatch rate | df | logLik | AICc | ΔAICc | Weight |
|---|---|---|---|---|---|---|---|
| Trento | CI | test vs control crosses | 4 | -587.70 | 1183.64 | 0.00 | 0.697 |
|  | Saturated | one estimate per cross type | 6 | -586.39 | 1185.31 | 1.67 | 0.303 |
|  | Null | intercept only | 3 | -624.42 | 1254.98 | 71.33 | 0.000 |
| Bal | Saturated | one estimate per cross type | 6 | -592.86 | 1198.25 | 0.00 | 0.771 |
|  | CI | test vs control crosses | 4 | -596.21 | 1200.68 | 2.43 | 0.229 |
|  | Null | intercept only | 3 | -651.45 | 1309.05 | 110.81 | 0.000 |
| Rennes | Saturated | one estimate per cross type | 6 | -343.91 | 700.70 | 0.00 | 0.793 |
|  | CI | test vs control crosses | 4 | -347.81 | 704.03 | 3.34 | 0.150 |
|  | Null | intercept only | 3 | -349.85 | 705.95 | 5.26 | 0.057 |
| Par6 | Null | intercept only | 3 | -187.54 | 381.49 | 0.00 | 0.546 |
|  | CI | test vs control crosses | 4 | -186.94 | 382.57 | 1.08 | 0.318 |
|  | Saturated | one estimate per cross type | 6 | -185.37 | 384.26 | 2.77 | 0.136 |

**Table S2b.** Mean hatch rates of the four cross types used to assay cytoplasmic incompatibility in four European *D. suzukii* lines. Hatch rates were estimated with a zero-inflated beta-binomial model including the effect of cross type, with a constant zero-inflation and a common dispersion parameter. Limits are 95% confidence limits obtained by parametric bootstrap with 500 iterations. Crosses are given as female × male. W+ denotes Wolbachia-infected, W-uninfected. Am is the uninfected tester line; the Am × Am cross is the common uninfected control (control cross 1).

| Line | Cross (♀ × ♂) | Cross type | n | Mean hatch rate | Lower 95% CI | Upper 95% CI |
|---|---|---|---|---|---|---|
| Trento | Am × Am | Control cross 1 W- x W- | 59 | 0.749 | 0.704 | 0.790 |
|  | Am × Trento | Test cross W- x W+ | 41 | 0.399 | 0.345 | 0.458 |
|  | Trento × Am | Control cross 2 W+ x W- | 34 | 0.704 | 0.642 | 0.765 |
|  | Trento × Trento | Control cross 3 W+ x W+ | 34 | 0.701 | 0.639 | 0.757 |
| Bal | Am × Am | Control cross 1 W- x W- | 59 | 0.732 | 0.680 | 0.783 |
|  | Am × Bal | Test cross W- x W+ | 46 | 0.246 | 0.188 | 0.313 |
|  | Bal × Am | Control cross 2 W+ x W- | 30 | 0.827 | 0.769 | 0.880 |
|  | Bal × Bal | Control cross 3 W+ x W+ | 31 | 0.774 | 0.703 | 0.834 |
| Rennes | Am × Am | Control cross 1 W- x W- | 33 | 0.756 | 0.687 | 0.816 |
|  | Am × Rennes | Test cross W- x W+ | 29 | 0.665 | 0.596 | 0.737 |
|  | Rennes × Am | Control cross 2 W+ x W- | 20 | 0.814 | 0.738 | 0.876 |
|  | Rennes × Rennes | Control cross 3 W+ x W+ | 20 | 0.663 | 0.584 | 0.749 |
| Par6 | Am × Am | Control cross 1 W- x W- | 21 | 0.732 | 0.605 | 0.822 |
|  | Am × Par6 | Test cross W- x W+ | 13 | 0.642 | 0.494 | 0.762 |
|  | Par6 × Am | Control cross 2 W+ x W- | 14 | 0.788 | 0.677 | 0.875 |
|  | Par6 × Par6 | Control cross 3 W+ x W+ | 14 | 0.628 | 0.483 | 0.760 |

**Table S2c.** Cytoplasmic incompatibility estimation for four European *D. suzukii* lines. 95% confidence intervals (CI) were obtained by parametric bootstrap with 500 iterations. Test crosses with the uninfected tester line Am are given for comparison as positive controls.

| **Line** | **Test cross**<br>**(♀ × ♂)** | **Control cross**<br>**(♀ × ♂)** | **Control type** | **CI strength** | **Lower 95% CI** | **Upper 95% CI** |
|---|---|---|---|---|---|---|
| Trento | Am × Trento | Am × Am | Control cross 1 (W− × W−) | 0.467 | 0.383 | 0.544 |
|  |  | Trento × Am | Control cross 2 (W+ × W−) | 0.433 | 0.335 | 0.520 |
|  |  | Trento × Trento | Control cross 3 (W+ × W+) | 0.430 | 0.330 | 0.520 |
| Bal | Am × Bal | Am × Am | Control cross 1 (W− × W−) | 0.665 | 0.576 | 0.748 |
|  |  | Bal × Am | Control cross 2 (W+ × W−) | 0.703 | 0.626 | 0.774 |
|  |  | Bal × Bal | Control cross 3 (W+ × W+) | 0.683 | 0.591 | 0.757 |
| Rennes | Am × Rennes | Am × Am | Control cross 1 (W− × W−) | 0.121 | 0.004 | 0.227 |
|  |  | Rennes × Am | Control cross 2 (W+ × W−) | 0.183 | 0.069 | 0.276 |
|  |  | Rennes × Rennes | Control cross 3 (W+ × W+) | 0.000 | 0.000 | 0.154 |
| Par6 | Am × Par6 | Am × Am | Control cross 1 (W− × W−) | 0.122 | 0.000 | 0.329 |
|  |  | Par6 × Am | Control cross 2 (W+ × W−) | 0.185 | 0.000 | 0.381 |
|  |  | Par6 × Par6 | Control cross 3 (W+ × W+) | 0.000 | 0.000 | 0.262 |

**Table S3a.**
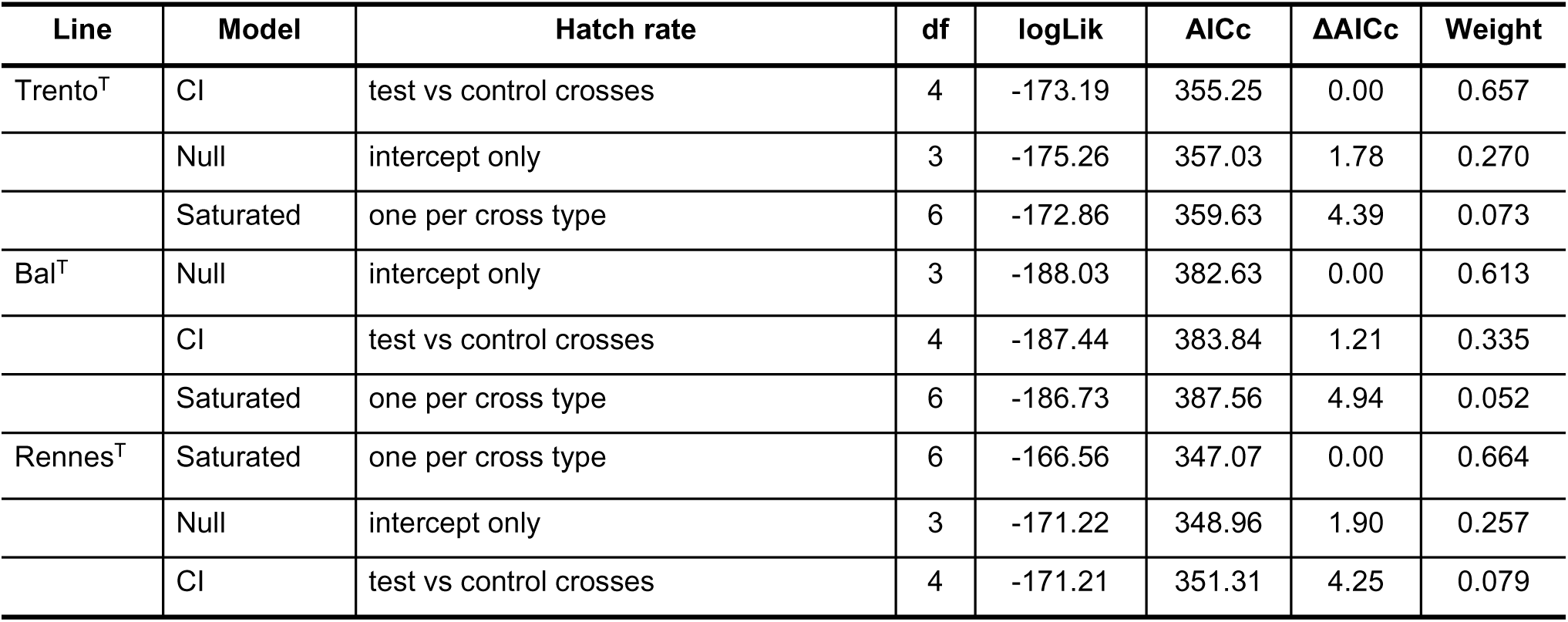
Results of the AICc model selection for tetracycline-treated lines of *D. suzukii*. Models are ranked from lowest to highest AICc values, and the first model listed for each strain corresponds to the best-supported model. All models included the identity of the mating pair as a random effect.

**Table S3b.** Mean hatch rates for tetracycline-treated lines of *D. suzukii*. Hatch rates were estimated with a zero-inflated beta-binomial model including the effect of cross type, with a constant zero-inflation and a common dispersion parameter. Limits are 95% confidence limits obtained by parametric bootstrap with 500 iterations. Crosses are given as female × male. Each line pair was analysed in a separate model.

| **Line** | **Cross (♀ × ♂)** | **Cross type** | **n** | **Mean hatch rate** | **Lower 95% CI** | **Upper 95% CI** |
|---|---|---|---|---|---|---|
| Trento<sup>T</sup> | Am × Am | Control cross 1 (W− × W−) | 12 | 0.752 | 0.651 | 0.837 |
|  | Am × Trento<sup>T</sup> | Test cross (W− × W+) | 9 | 0.648 | 0.507 | 0.760 |
|  | Trento<sup>T</sup> × Am | Control cross 2 (W+ × W−) | 15 | 0.799 | 0.714 | 0.881 |
|  | Trento<sup>T</sup> × Trento<sup>T</sup> | Control cross 3 (W+ × W+) | 15 | 0.772 | 0.657 | 0.859 |
| Bal<sup>T</sup> | Am × Am | Control cross 1 (W− × W−) | 12 | 0.737 | 0.623 | 0.833 |
|  | Am × Bal<sup>T</sup> | Test cross (W− × W+) | 9 | 0.692 | 0.541 | 0.800 |
|  | Bal<sup>T</sup> × Am | Control cross 2 (W+ × W−) | 12 | 0.811 | 0.714 | 0.895 |
|  | Bal<sup>T</sup> × Bal<sup>T</sup> | Control cross 3 (W+ × W+) | 14 | 0.751 | 0.649 | 0.833 |
| Rennes<sup>T</sup> | Am × Am | Control cross 1 (W− × W−) | 12 | 0.756 | 0.673 | 0.831 |
|  | Am × Rennes<sup>T</sup> | Test cross (W− × W+) | 10 | 0.768 | 0.682 | 0.837 |
|  | Rennes<sup>T</sup> × Am | Control cross 2 (W+ × W−) | 15 | 0.845 | 0.787 | 0.898 |
|  | Rennes<sup>T</sup> × Rennes<sup>T</sup> | Control cross 3 (W+ × W+) | 13 | 0.692 | 0.609 | 0.774 |

**Table S3c.** Cytoplasmic incompatibility estimation for tetracycline-treated lines of *D. suzukii*. 95% confidence intervals (CI) were obtained by parametric bootstrap with 500 iterations. Test crosses with the uninfected tester line Am are given for comparison as positive controls.

| **Line** | **Test cross**<br>**(♀ × ♂)** | **Control cross**<br>**(♀ × ♂)** | **Control type** | **CI strength** | **Lower 95% CI** | **Upper 95% CI** |
|---|---|---|---|---|---|---|
| Trento<sup>T</sup> | Am × Trento<sup>T</sup> | Am × Am | Control cross 1 (W− × W−) | 0.139 | 0.000 | 0.327 |
|  |  | Trento<sup>T</sup> × Am | Control cross 2 (W+ × W−) | 0.189 | 0.012 | 0.382 |
|  |  | Trento<sup>T</sup> Trento<sup>T</sup> | Control cross 3 (W+ × W+) | 0.161 | 0.000 | 0.358 |
| Bal<sup>T</sup> | Am × Bal<sup>T</sup> | Am × Am | Control cross 1 (W− × W−) | 0.061 | 0.000 | 0.260 |
|  |  | Bal<sup>T</sup> × Am | Control cross 2 (W+ × W−) | 0.147 | 0.000 | 0.346 |
|  |  | Bal<sup>T</sup> × Bal<sup>T</sup> | Control cross 3 (W+ × W+) | 0.080 | 0.000 | 0.274 |
| Rennes<sup>T</sup> | Am × Rennes<sup>T</sup> | Am × Am | Control cross 1 (W− × W−) | 0.000 | 0.000 | 0.123 |
|  |  | Rennes<sup>T</sup> × Am | Control cross 2 (W+ × W−) | 0.091 | 0.000 | 0.207 |
|  |  | Rennes<sup>T</sup> × Rennes<sup>T</sup> | Control cross 3 (W+ × W+) | 0.000 | 0.000 | 0.052 |

**Table S4.** Pairwise comparisons of *Wolbachia* titer in testes among four European *D. suzukii* lines. Wolbachia titers were log_₁₀_-transformed before analysis; lines were compared by ANOVA (F_₃_,_₁₆_ = 6.17, P = 0.0055) followed by pairwise contrasts. Ratios of titers and their confidence intervals were obtained by back-transformation from the log_₁₀_ scale; a ratio of 1 indicates equal titers. P values and confidence intervals are adjusted for multiple comparisons by the Tukey method for a family of four estimates. n = 5 samples of 5 pairs of testes per line.

| **Lines** | **Ratio of titers** | **SE** | **df** | **t ratio** | **Lower 95% CI** | **Upper 95% CI** | **P value** |
|---|---|---|---|---|---|---|---|
| Trento / Bal | 5.606 | 2.685 | 16 | 3.599 | 1.424 | 22.070 | 0.0116 |
| Trento / Par6 | 5.370 | 2.572 | 16 | 3.509 | 1.364 | 21.140 | 0.0139 |
| Trento / Rennes | 1.883 | 0.902 | 16 | 1.321 | 0.478 | 7.410 | 0.5634 |
| Bal / Par6 | 0.958 | 0.459 | 16 | -0.090 | 0.243 | 3.770 | 0.9997 |
| Bal / Rennes | 0.336 | 0.161 | 16 | -2.278 | 0.085 | 1.320 | 0.1450 |
| Par6 / Rennes | 0.351 | 0.168 | 16 | -2.188 | 0.089 | 1.380 | 0.1688 |

**Table S5a.** Results of the AICc model selection for hatch rate in crosses between *Wolbachia*-infected *D. suzukii* lines (rescue crosses) Models are ranked from lowest to highest AICc values, and the first model listed for each strain corresponds to the best-supported model.

| **Male line** | **Model** | **Hatch rate** | **df** | **logLik** | **AICc** | **ΔAICc** | **Weight** |
|---|---|---|---|---|---|---|---|
| Trento | Saturated | one estimate per cross | 5 | -221.42 | 454.00 | 0.00 | 0.596 |
|  | CI | one estimate for CI vs. rescue crosses | 4 | -223.02 | 454.80 | 0.80 | 0.400 |
|  | Null | intercept only | 3 | -228.84 | 464.13 | 10.13 | 0.004 |
| Bal | Saturated | one estimate per cross | 6 | -173.62 | 361.19 | 0.00 | 0.855 |
|  | CI | one estimate for CI vs. rescue crosses | 4 | -177.93 | 364.74 | 3.55 | 0.145 |
|  | Null | intercept only | 3 | -187.55 | 381.62 | 20.43 | 0.000 |

**Table S5b.** Mean hatch rates of crosses between *Wolbachia*-infected *D. suzukii* lines (rescue crosses) Hatch rates were estimated separately for each male line with a zero-inflated beta-binomial model including the effect of cross type, with a constant zero-inflation and a common dispersion parameter. Limits are 95% confidence limits obtained by parametric bootstrap with 500 iterations. Crosses are given as female × male.

| **Male line** | **Cross (♀ × ♂)** | **Cross type** | **n** | **Mean hatch rate** | **Lower 95% CI** | **Upper 95% CI** |
|---|---|---|---|---|---|---|
| Trento | Am × Trento | CI cross W- x W+ | 20 | 0.463 | 0.375 | 0.555 |
|  | Trento × Trento | Rescue cross W+ x W+ | 27 | 0.630 | 0.566 | 0.698 |
|  | Bal × Trento | Rescue cross W+ x W+ | 11 | 0.744 | 0.639 | 0.833 |
| Bal | Am × Bal | CI cross W- x W+ | 19 | 0.246 | 0.137 | 0.351 |
|  | Bal × Bal | Rescue cross W+ x W+ | 11 | 0.835 | 0.718 | 0.916 |
|  | Trento × Bal | Rescue cross W+ x W+ | 10 | 0.644 | 0.485 | 0.775 |
|  | Par6 × Bal | Rescue cross W+ x W+ | 10 | 0.511 | 0.344 | 0.675 |

**Table S5c.** Reduction in hatch rate of CI crosses with uninfected Am females relative to rescue crosses with infected females. CI strength is the relative reduction in hatch rate of the test cross (Am females) compared with each control cross (infected females), truncated at zero. Values above zero indicate a higher hatch rate with infected females, as expected under rescue. Estimates were derived from the saturated model (Table S5a); 95% confidence intervals (CI) were obtained by parametric bootstrap.

| Male line | CI cross<br>(♀ × ♂) | Rescue cross<br>(♀ × ♂) | Control female line | CI strength | Lower 95% CI | Upper 95% CI |
|---|---|---|---|---|---|---|
| Trento | Am × Trento | Trento × Trento | Same line (W+ × W+) | 0.266 | 0.078 | 0.423 |
|  |  | Bal × Trento | Other line (W+ × W+) | 0.378 | 0.184 | 0.515 |
| Bal | Am × Bal | Bal × Bal | Same line (W+ × W+) | 0.705 | 0.572 | 0.840 |
|  |  | Trento × Bal | Other line (W+ × W+) | 0.617 | 0.415 | 0.796 |
|  |  | Par6 × Bal | Other line (W+ × W+) | 0.517 | 0.177 | 0.738 |

**Table S6a.** Results of the AICc model selection for hatch rate after introgression of four *w*Suz isolates into the Am nuclear background. Models are ranked from lowest to highest AICc values, and the first model listed for each strain corresponds to the best-supported model. All models included the identity of the mating pair as a random effect. Introgressed lines carry the Wolbachia and cytoplasm of the donor line in the nuclear background of the uninfected Am line after 7 generations of backcrossing; Am is the uninfected recipient line.

| Line | Model | Hatch rate | df | logLik | AICc | ΔAICc | Weight |
|---|---|---|---|---|---|---|---|
| Am<sup>Trento</sup> | CI | test vs control crosses | 4 | -284.45 | 577.39 | 0.00 | 0.434 |
|  | Null | intercept only | 3 | -285.66 | 577.61 | 0.23 | 0.388 |
|  | Saturated | one per cross type | 6 | -283.06 | 579.16 | 1.78 | 0.179 |
| Am<sup>Bal</sup> | Null | intercept only | 3 | -241.50 | 489.31 | 0.00 | 0.631 |
|  | CI | test vs control crosses | 4 | -241.12 | 490.78 | 1.47 | 0.303 |
|  | Saturated | one per cross type | 6 | -240.33 | 493.82 | 4.51 | 0.066 |
| Am<sup>Rennes</sup> | Null | intercept only | 3 | -246.06 | 498.42 | 0.00 | 0.609 |
|  | CI | test vs control crosses | 4 | -245.52 | 499.52 | 1.11 | 0.350 |
|  | Saturated | one per cross type | 6 | -245.39 | 503.83 | 5.42 | 0.041 |
| Am<sup>Par6</sup> | Null | intercept only | 3 | -204.89 | 416.10 | 0.00 | 0.677 |
|  | CI | test vs control crosses | 4 | -204.73 | 418.03 | 1.93 | 0.259 |
|  | Saturated | one per cross type | 6 | -203.81 | 420.83 | 4.73 | 0.064 |

**Table S6b.**
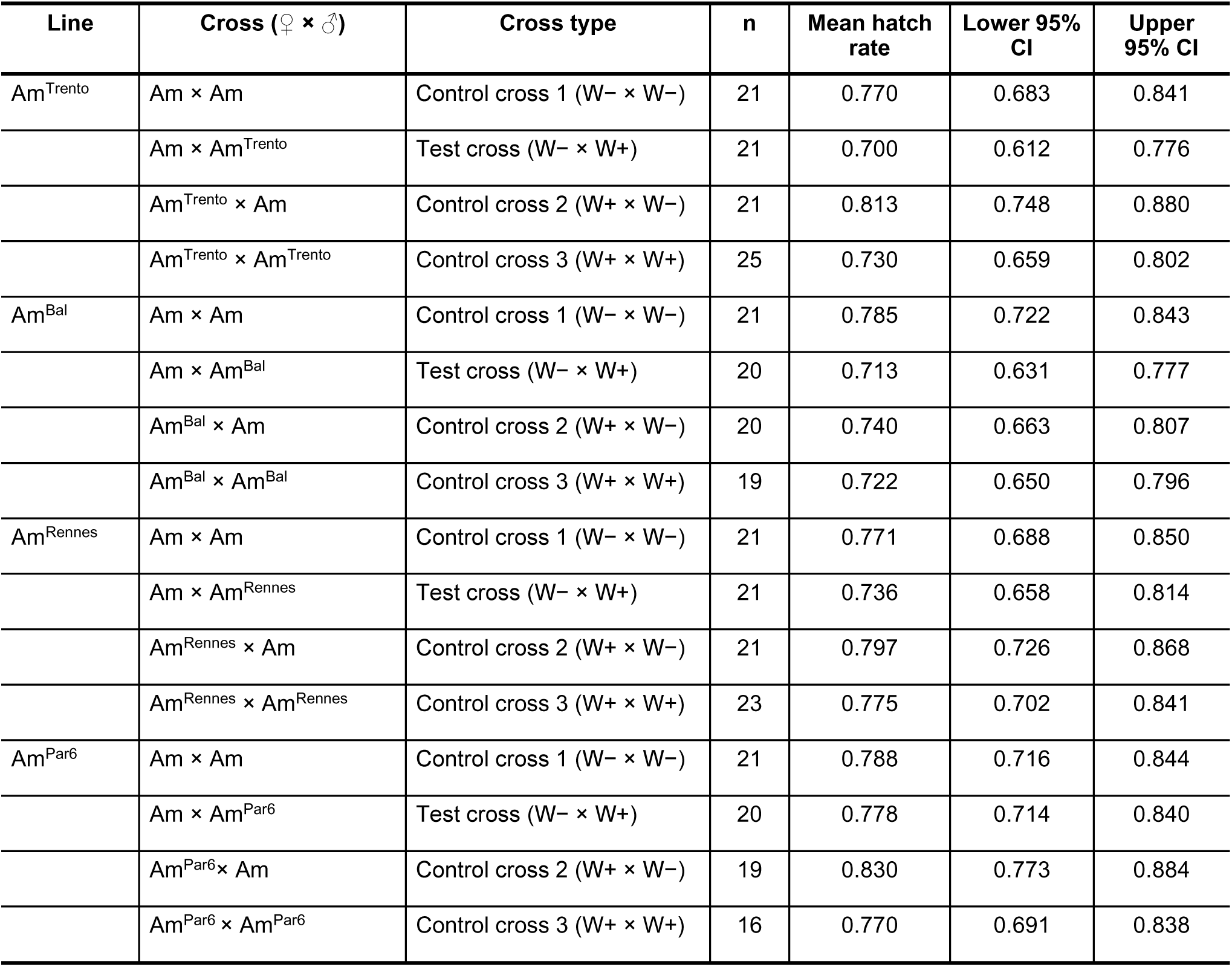
Mean hatch rates for introgressed lines of *D. suzukii*. Hatch rates were estimated with a zero-inflated beta-binomial model including the effect of cross type, with a constant zero-inflation and a common dispersion parameter. Limits are 95% confidence limits obtained by parametric bootstrap with 500 iterations. Crosses are given as female × male. Each line pair was analysed in a separate model.

| Line | Cross (♀ × ♂) | Cross type | n | Mean hatch rate | Lower 95% CI | Upper 95% CI |
|---|---|---|---|---|---|---|
| Am<sup>Trento</sup> | Am × Am | Control cross 1 (W− × W−) | 21 | 0.770 | 0.683 | 0.841 |
|  | Am × Am<sup>Trento</sup> | Test cross (W− × W+) | 21 | 0.700 | 0.612 | 0.776 |
|  | Am<sup>Trento</sup> × Am | Control cross 2 (W+ × W−) | 21 | 0.813 | 0.748 | 0.880 |
|  | Am<sup>Trento</sup> × Am<sup>Trento</sup> | Control cross 3 (W+ × W+) | 25 | 0.730 | 0.659 | 0.802 |
| Am<sup>Bal</sup> | Am × Am | Control cross 1 (W− × W−) | 21 | 0.785 | 0.722 | 0.843 |
|  | Am × Am<sup>Bal</sup> | Test cross (W− × W+) | 20 | 0.713 | 0.631 | 0.777 |
|  | Am<sup>Bal</sup> × Am | Control cross 2 (W+ × W−) | 20 | 0.740 | 0.663 | 0.807 |
|  | Am<sup>Bal</sup> × Am<sup>Bal</sup> | Control cross 3 (W+ × W+) | 19 | 0.722 | 0.650 | 0.796 |
| Am<sup>Rennes</sup> | Am × Am | Control cross 1 (W− × W−) | 21 | 0.771 | 0.688 | 0.850 |
|  | Am × Am<sup>Rennes</sup> | Test cross (W− × W+) | 21 | 0.736 | 0.658 | 0.814 |
|  | Am<sup>Rennes</sup> × Am | Control cross 2 (W+ × W−) | 21 | 0.797 | 0.726 | 0.868 |
|  | Am<sup>Rennes</sup> × Am<sup>Rennes</sup> | Control cross 3 (W+ × W+) | 23 | 0.775 | 0.702 | 0.841 |
| Am<sup>Par6</sup> | Am × Am | Control cross 1 (W− × W−) | 21 | 0.788 | 0.716 | 0.844 |
|  | Am × Am<sup>Par6</sup> | Test cross (W− × W+) | 20 | 0.778 | 0.714 | 0.840 |
|  | Am<sup>Par6</sup>× Am | Control cross 2 (W+ × W−) | 19 | 0.830 | 0.773 | 0.884 |
|  | Am<sup>Par6</sup> × Am<sup>Par6</sup> | Control cross 3 (W+ × W+) | 16 | 0.770 | 0.691 | 0.838 |

**Table S6c.** Cytoplasmic incompatibility estimation for introgressed lines of *D. suzukii*. 95% confidence intervals (CI) were obtained by parametric bootstrap with 500 iterations. Test crosses with the uninfected tester line Am are given for comparison as positive controls.

| Line | Test cross<br>(♀ × ♂) | Control cross<br>(♀ × ♂) | Control type | CI strength | Lower 95% CI | Upper 95% CI |
|---|---|---|---|---|---|---|
| Am<sup>Trento</sup> | Am × Am<sup>Trento</sup> | Am × Am | Control cross 1 (W− × W−) | 0.092 | 0.000 | 0.227 |
|  |  | Am<sup>Trento</sup> × Am | Control cross 2 (W+ × W−) | 0.139 | 0.027 | 0.261 |
|  |  | Am<sup>Trento</sup> × Am<sup>Trento</sup> | Control cross 3 (W+ × W+) | 0.042 | 0.000 | 0.185 |
| Am<sup>Bal</sup> | Am × Am<sup>Bal</sup> | Am × Am | Control cross 1 (W− × W−) | 0.092 | 0.000 | 0.205 |
|  |  | Am<sup>Bal</sup> × Am | Control cross 2 (W+ × W−) | 0.037 | 0.000 | 0.165 |
|  |  | Am<sup>Bal</sup> × Am<sup>Bal</sup> | Control cross 3 (W+ × W+) | 0.013 | 0.000 | 0.147 |
| Am<sup>Rennes</sup> | Am × Am<sup>Rennes</sup> | Am × Am | Control cross 1 (W− × W−) | 0.045 | 0.000 | 0.177 |
|  |  | Am<sup>Rennes</sup> × Am | Control cross 2 (W+ × W−) | 0.076 | 0.000 | 0.201 |
|  |  | Am<sup>Rennes</sup> × Am<sup>Rennes</sup> | Control cross 3 (W+ × W+) | 0.050 | 0.000 | 0.163 |
| Am<sup>Par6</sup> | Am × Am<sup>Par6</sup> | Am × Am | Control cross 1 (W− × W−) | 0.012 | 0.000 | 0.113 |
|  |  | Am<sup>Par6</sup>× Am | Control cross 2 (W+ × W−) | 0.063 | 0.000 | 0.165 |
|  |  | Am<sup>Par6</sup> × Am<sup>Par6</sup> | Control cross 3 (W+ × W+) | 0.000 | 0.000 | 0.103 |

**Table S7a.**
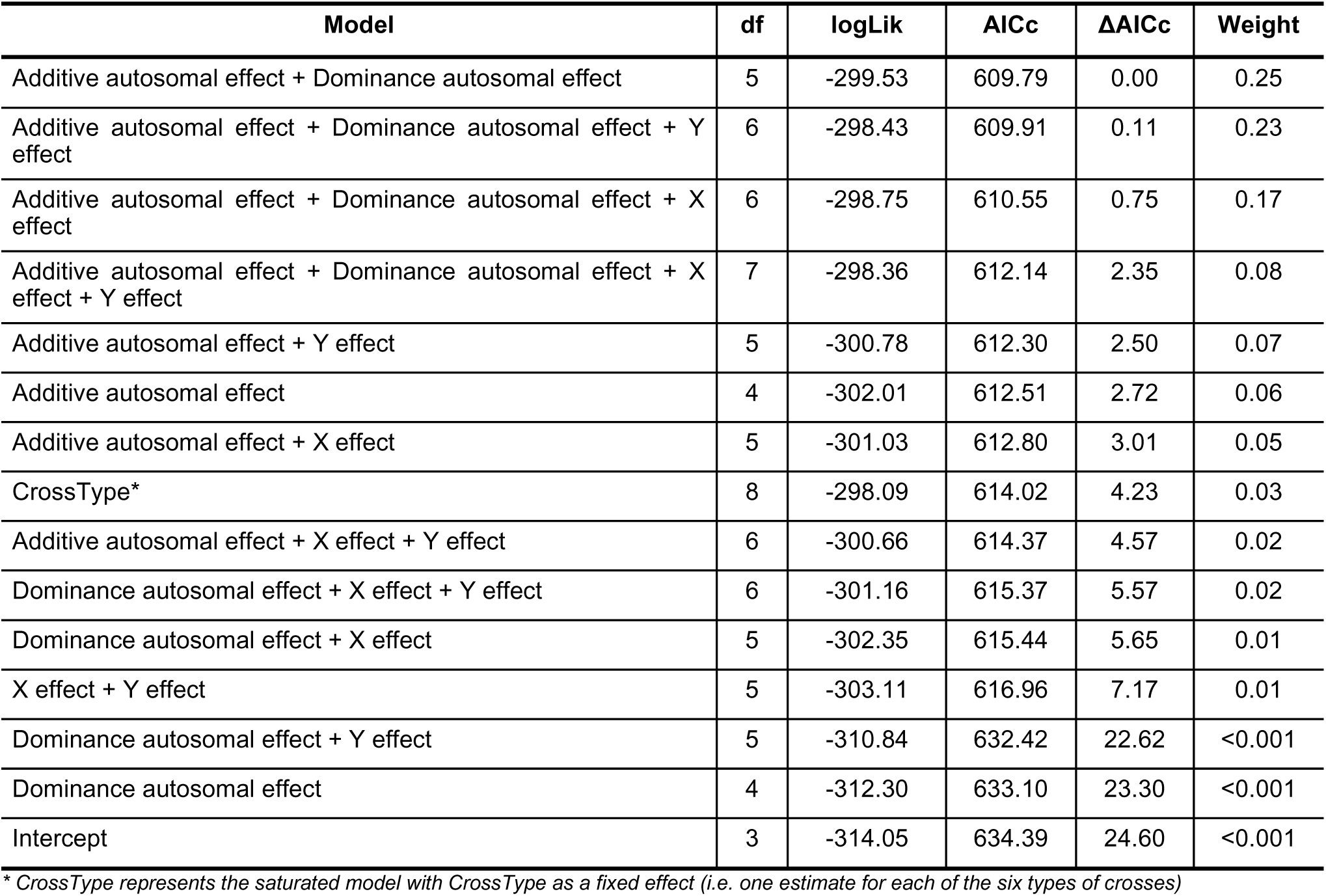
AICc model selection for the genetic architecture of CI suppression. Zero-inflated beta-binomial Generalized Linear Mixed Models combining additive autosomal, dominance autosomal, X-linked and Y-linked effects, fitted to hatch rates in test crosses with parental, F1 and F2 males Models are ranked by increasing AICc, from best to least supported. All models included a constant zero-inflation probability. df, number of estimated parameters; logLik, log-likelihood; ΔAICc, difference in AICc from the best model; weight, Akaike’s weight.

| **Model** | **df** | **logLik** | **AICc** | **ΔAICc** | **Weight** |
|---|---|---|---|---|---|
| Additive autosomal effect + Dominance autosomal effect | 5 | -299.53 | 609.79 | 0.00 | 0.25 |
| Additive autosomal effect + Dominance autosomal effect + Y effect | 6 | -298.43 | 609.91 | 0.11 | 0.23 |
| Additive autosomal effect + Dominance autosomal effect + X effect | 6 | -298.75 | 610.55 | 0.75 | 0.17 |
| Additive autosomal effect + Dominance autosomal effect + X effect + Y effect | 7 | -298.36 | 612.14 | 2.35 | 0.08 |
| Additive autosomal effect + Y effect | 5 | -300.78 | 612.30 | 2.50 | 0.07 |
| Additive autosomal effect | 4 | -302.01 | 612.51 | 2.72 | 0.06 |
| Additive autosomal effect + X effect | 5 | -301.03 | 612.80 | 3.01 | 0.05 |
| CrossType* | 8 | -298.09 | 614.02 | 4.23 | 0.03 |
| Additive autosomal effect + X effect + Y effect | 6 | -300.66 | 614.37 | 4.57 | 0.02 |
| Dominance autosomal effect + X effect + Y effect | 6 | -301.16 | 615.37 | 5.57 | 0.02 |
| Dominance autosomal effect + X effect | 5 | -302.35 | 615.44 | 5.65 | 0.01 |
| X effect + Y effect | 5 | -303.11 | 616.96 | 7.17 | 0.01 |
| Dominance autosomal effect + Y effect | 5 | -310.84 | 632.42 | 22.62 | <0.001 |
| Dominance autosomal effect | 4 | -312.30 | 633.10 | 23.30 | <0.001 |
| Intercept | 3 | -314.05 | 634.39 | 24.60 | <0.001 |

*\* CrossType represents the saturated model with CrossType as a fixed effect (i.e. one estimate for each of the six types of crosses)*

**Table S7b.** Genetic effects on hatch rate estimated from the full generation means model. Parameter estimates (logit scale) from a zero-inflated beta-binomial Generalized Linear Mixed Model including additive autosomal (a), dominance autosomal (d), X-linked (x) and Y-linked (y) effects, fitted to test crosses between Am females and parental (Bal, Par6), F1 and F2 males. 95% confidence interval (CI) were computed using parametric bootstrap with 500 iterations.

| Effect | Estimate | Lower 95% CI | Upper 95% CI |
|---|---|---|---|
| **Zero-inflation part** |  |  |  |
| (Intercept) | -2.726 | -4.311 | -1.957 |
| **Conditional (beta-binomial) part** |  |  |  |
| (Intercept) | -0.139 | -0.486 | 0.163 |
| Additive effect | 1.021 | 0.188 | 1.944 |
| Dominance effect | 0.571 | 0.089 | 1.123 |
| X effect | 0.093 | -0.447 | 0.622 |
| Y effect | -0.147 | -0.466 | 0.158 |

**Table S7c.** Estimated hatch rates for each type of cross. Hatch rates were estimated from a zero-inflated beta-binomial Generalized Linear Mixed Model with cross type as a fixed effect and couple as a random intercept. The expected hatch rate was calculated as (1 − π)μ, where π is the probability of a structural zero (a couple with no offspring larvae) and μ is the mean probability in the beta-binomial component. Estimates are for a typical couple (random effect set to zero). 95% confidence interval (CI) were computed using parametric bootstrap with 500 iterations.

| **Type of cross (♀ × ♂)** | **n** | **Hatch rate** | **Lower 95% CI** | **Upper 95% CI** |
|---|---|---|---|---|
| AmxAm | 16 | 0.654 | 0.547 | 0.746 |
| AmxBal | 17 | 0.253 | 0.168 | 0.349 |
| AmxF1 BalxPar6 | 12 | 0.566 | 0.431 | 0.687 |
| AmxF1 Par6xBal | 12 | 0.681 | 0.559 | 0.780 |
| AmxF2 (BalxPar6)x(BalxPar6) | 16 | 0.475 | 0.370 | 0.578 |
| AmxF2 (Par6xBal)x(Par6xBal) | 14 | 0.551 | 0.446 | 0.654 |
| AmxPar6 | 16 | 0.712 | 0.622 | 0.801 |
| BalxAm | 20 | 0.748 | 0.677 | 0.817 |
| BalxBal | 20 | 0.726 | 0.644 | 0.805 |
| F1 BalxPar6xAm | 12 | 0.760 | 0.649 | 0.849 |
| F1 BalxPar6xF1 BalxPar6 | 12 | 0.732 | 0.620 | 0.826 |
| F1 Par6xBalxAm | 8 | 0.786 | 0.670 | 0.877 |
| F1 Par6xBalxF1 Par6xBal | 8 | 0.759 | 0.631 | 0.857 |
| F2 (BalxPar6)x(BalxPar6)xAm | 14 | 0.843 | 0.760 | 0.909 |
| F2 (BalxPar6)x(BalxPar6)xF2 (BalxPar6)x(BalxPar6) | 11 | 0.734 | 0.619 | 0.838 |
| F2 (Par6xBal)x(Par6xBal)xAm | 14 | 0.716 | 0.615 | 0.811 |
| F2 (Par6xBal)x(Par6xBal)xF2 (Par6xBal)x(Par6xBal) | 8 | 0.729 | 0.611 | 0.838 |
| Par6xAm | 18 | 0.794 | 0.715 | 0.858 |
| Par6xBal | 6 | 0.832 | 0.700 | 0.928 |
| Par6xPar6 | 18 | 0.445 | 0.347 | 0.542 |

**Table S8a.** Results of the AICc model selection for variation in CI levels among uninfected female genotypes. Models are ranked from lowest to highest AICc values, and the first model listed for each strain corresponds to the best-supported model.

| **Line** | **Model** | **Hatch rate** | **df** | **logLik** | **AICc** | **ΔAICc** | **Weight** |
|---|---|---|---|---|---|---|---|
| Bal | Null | intercept only | 3 | -159.805 | 326.168 | 0 | 0.867 |
|  | Female genotype | one estimate per female genotype | 5 | -159.231 | 329.925 | 3.756 | 0.133 |
| Rennes | Null | intercept only | 3 | -186.674 | 379.892 | 0 | 0.724 |
|  | Female genotype | one estimate per female genotype | 4 | -186.444 | 381.819 | 1.926 | 0.276 |
| Par6 | Null | intercept only | 3 | -177.458 | 361.475 | 0 | 0.768 |
|  | Female genotype | one estimate per female genotype | 4 | -177.457 | 363.866 | 2.391 | 0.232 |

**Table S8b.**
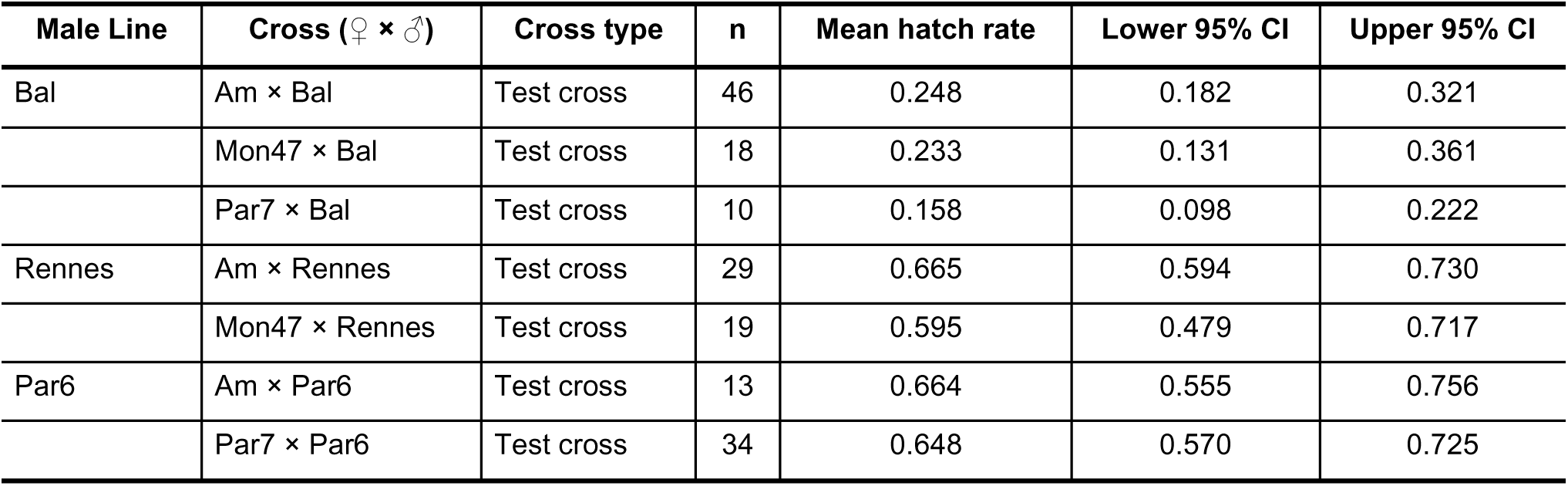
Mean hatch rates of crosses between *Wolbachia*-infected males lines and different female genotypes. Hatch rates were estimated with a zero-inflated beta-binomial model including the effect of cross type, with a constant zero-inflation and a common dispersion parameter. Limits are 95% confidence limits obtained by parametric bootstrap with 500 iterations. Crosses are given as female × male. Each line pair was analysed in a separate model. Am, Mon47 and Par7 are uninfected female lines.

**Table S9a.** Selection between zero-inflated beta-binomial models of hatch rate with and without the effect of male age. Models differ only in the structure of the hatch-rate part; all include a constant zero-inflation term and a common dispersion parameter. No age: hatch rate differs among male lines only. Age: male line plus a male age effect common to all lines. Models are ordered by AICc within each analysis; ΔAICc is the difference from the best-supported model of that analysis. ΔAICc < 2 is not taken as evidence for an effect.

| **Analysis** | **Model** | **Fixed effects** | **df** | **logLik** | **AICc** | **ΔAICc** | **Weight** |
|---|---|---|---|---|---|---|---|
| Test crosses, four lines | No age | male line | 6 | -472.462 | 957.612 | 0.000 | 0.748 |
|  | Age | male line + male age | 7 | -472.432 | 959.790 | 2.178 | 0.252 |
| Test crosses, Bal and Trento | No age | male line | 4 | -321.556 | 651.600 | 0.000 | 0.751 |
|  | Age | male line + male age | 5 | -321.535 | 653.811 | 2.211 | 0.249 |
| Control cross 3, four lines | No age | male line | 6 | -462.155 | 937.010 | 0.000 | 0.721 |
|  | Age | male line + male age | 7 | -461.985 | 938.912 | 1.902 | 0.279 |

**Table S9b.** Mean hatch rate by male line and male age in test crosses and control crosses 3 with the same male. *Hatch rates* were estimated with zero-inflated beta-binomial models including the effects of male line and male age, with a constant zero-inflation and a common dispersion parameter, fitted separately to test crosses and to control crosses 3. Estimates were taken from the model including male age, which assumes the same age effect in all lines. 95% confidence intervals (CI) were obtained by parametric bootstrap (500 replicates; percentile intervals). Crosses are given as female × male. W+ denotes Wolbachia-infected, W− uninfected; Am is the uninfected tester line. Male age is given in days post-eclosion. n is the number of couples. Only line × age combinations that were tested are shown.

| Line | Cross (♀ × ♂) | Cross type | Male age (days) | n | Mean hatch rate | Lower 95% CI | Upper 95% CI |
|---|---|---|---|---|---|---|---|
| Trento | Am × Trento | Test cross W- x W+ | 2.0 | 12 | 0.375 | 0.283 | 0.467 |
|  |  |  | 2.5 | 10 | 0.381 | 0.310 | 0.460 |
|  |  |  | 3.0 | 19 | 0.388 | 0.301 | 0.483 |
|  | Trento × Trento | Control cross 3 W+ x W+ | 2.0 | 16 | 0.593 | 0.507 | 0.673 |
|  |  |  | 2.5 | 5 | 0.607 | 0.536 | 0.676 |
|  |  |  | 3.0 | 22 | 0.621 | 0.527 | 0.699 |
| Bal | Am × Bal | Test cross W- x W+ | 2.0 | 7 | 0.225 | 0.148 | 0.306 |
|  |  |  | 2.5 | 16 | 0.230 | 0.176 | 0.291 |
|  |  |  | 3.0 | 23 | 0.235 | 0.180 | 0.295 |
|  | Bal × Bal | Control cross 3 W+ x W+ | 2.0 | 2 | 0.696 | 0.566 | 0.782 |
|  |  |  | 2.5 | 2 | 0.707 | 0.614 | 0.775 |
|  |  |  | 3.0 | 27 | 0.719 | 0.641 | 0.785 |
| Rennes | Am × Rennes | Test cross W- x W+ | 2.0 | 6 | 0.623 | 0.514 | 0.727 |
|  |  |  | 2.5 | 5 | 0.629 | 0.548 | 0.709 |
|  |  |  | 3.0 | 18 | 0.636 | 0.547 | 0.721 |
|  | Rennes × Rennes | Control cross 3 W+ x W+ | 2.0 | 4 | 0.606 | 0.499 | 0.708 |
|  |  |  | 2.5 | 7 | 0.620 | 0.526 | 0.704 |
|  |  |  | 3.0 | 16 | 0.633 | 0.541 | 0.713 |
| Par6 | Am × Par6 | Test cross W- x W+ | 3.0 | 13 | 0.630 | 0.504 | 0.753 |
|  | Par6 × Par6 | Control cross 3 W+ x W+ | 2.0 | 8 | 0.529 | 0.420 | 0.635 |
|  |  |  | 2.5 | 4 | 0.544 | 0.452 | 0.632 |
|  |  |  | 3.0 | 14 | 0.558 | 0.466 | 0.652 |

**Table S9c.** Effect of male age on hatch rate. The age effect is given as the odds ratio of hatching per additional day of male age (exponentiated slope on the logit scale) in the model including male age (Table S9a); it is common to all lines of the analysis. Male age ranged from 2 to 3 days post-eclosion; lines tested at a single male age do not contribute to the estimate. 95% confidence intervals (CI) were obtained by parametric bootstrap (500 replicates; percentile intervals); the last column gives the number of converged bootstrap refits. n is the number of couples. A confidence interval that includes 1 means that no age effect was detected; it is not evidence that male age has no effect.

| **Analysis** | **Crosses (♀ × ♂)** | **n** | **Odds ratio per day** | **Lower 95% CI** | **Upper 95% CI** |
|---|---|---|---|---|---|
| Test crosses, four lines | W− × W+ | 129 | 1.061 | 0.690 | 1.714 |
| Test crosses, Bal and Trento | W− × W+ | 87 | 0.937 | 0.545 | 1.806 |
| Control cross 3, four lines | W+ × W+ | 127 | 1.139 | 0.735 | 1.718 |

**Table S10.** Whole-genome sequencing data used for the phylogenomic analyses of *Wolbachia* isolates. Sample accessions are given for the European Nucleotide Archive (ENA) or the Sequence Read Archive (SRA). Isolates sequenced in this study are indicated in the Reference column.

| Species | ID | Isolate | Sample accession | Location | Year | Reference | Comments |
|---|---|---|---|---|---|---|---|
| *D. suzukii* | Iso-FR-Par006 | Par6 | ERR18017645 | Paris, France | 2014 | This study |  |
| *D. suzukii* | Col-FR-Bal001 | Bal | ERR18017642 | Bellegarde, France | 2013 | This study |  |
| *D. suzukii* | Col-IT-Tre000 | Trento IAEA | ERR18017644 | Trento Province, Italy | Before 2013 | This study |  |
| *D. suzukii* | Col-FR-Ren000 | Rennes | ERR18017643 | Thorigné-Fouillard, France | 2017 | This study |  |
| *D. suzukii* | IT-Tre-Om300 | IT-Tre-Om300 | ERP001893 | Valsugana, Trento Province, Italy | Unknown | Ometto et al 2013 |  |
| *D. suzukii* | RW-2025a | RW-2025a | SRX27255168 | Yantai, China | 2018 | Wang et al. 2025 |  |
| *D. subpulchrella* | D-sub | D-sub | SRR29892916 | Narusawa-mura, Japan | 2016 | Muto et al 2018,<br>Camus et al 2025 | Mix of lines NAR01 and NAR02 |
| *D. subpulchrella* | Ko-Nam_NW2 | Ko-Nam_NW2 | SRX10342067 | Namwon, South Korea | 2016 | Lewald et al 2021;<br>Gautier 2023 | Korea-AA7 in Turelli 2018 |
| *D. subpulchrella* | CN-Kun_KM5 | CN-Kun_KM5 | SRX10342257 | Kunming, Yunnan, China | 2016 | Lewald et al 2021;<br>Gautier 2023 | China-AA27 in Turelli 2018 |
| *D. simulans* | US-CA_wRI_Dsim2 | Y14_29 | SRS2899949 | Yolo, California, USA | 2014 | Turelli et al 2018 |  |
| *D. simulans* | ZB_wRI_Dsim1 | NMB15_030 | SRS2899940 | Zambia | 2015 | Turelli et al 2018 |  |

**References**
Gautier, M. (2023). Efficient k-mer based curation of raw sequence data: application in *Drosophila suzukii*. Peer Community Journal, 3.
Lewald, K. M., Abrieux, A., Wilson, D. A., Lee, Y., Conner, W. R., Andreazza, F., ... & Chiu, J. C. (2021). Population genomics of *Drosophila suzukii* reveal longitudinal population structure and signals of migrations in and out of the continental United States. G3, 11(12), jkab343.
Muto, L., Kamimura, Y., Tanaka, K. M., & Takahashi, A. (2018). An innovative ovipositor for niche exploitation impacts genital coevolution between sexes in a fruit-damaging *Drosophila*. Proceedings of the Royal Society B: Biological Sciences, 285(1887), 20181635.
Ometto, L., Cestaro, A., Ramasamy, S., Grassi, A., Revadi, S., Siozios, S., ... & Rota-Stabelli, O. (2013). Linking genomics and ecology to investigate the complex evolution of an invasive *Drosophila* pest. Genome biology and evolution, 5(4), 745-757.
Turelli, M., Cooper, B. S., Richardson, K. M., Ginsberg, P. S., Peckenpaugh, B., Antelope, C. X., ... & Hoffmann, A. A. (2018). Rapid global spread of wRi-like *Wolbachia* across multiple *Drosophila*. Current Biology, 28(6), 963-971.

